# Robust High-Throughput Imaging Analysis with Wasserstein Geodesic Transformations

**DOI:** 10.1101/2025.10.01.679822

**Authors:** Gregory J. Hunt, Johann A. Gagnon-Bartsch

## Abstract

High-throughput cell imaging has been increasingly used in drug discovery to simultaneously profile the morphological response of cells to thousands of compounds using high-resolution microscopy and automated image analysis. Such experiments characterize thousands of image features in millions of cells across thousands of experimental conditions. Analytical difficulties arise with this scale of analysis as many features have distributions with extremely long tails, high skewness, remote outliers, and high-leverage points. This makes important signals difficult to find and means analyses are often sensitive to individual observations or features.

This work considers a recent high-quality Cell Painting dataset profiling compounds from the EU-OPENSCREEN consortium. The study perturbs HepG2 human liver cancer cells in order to morphologically profile cellular response to the compounds. Without adjustment, analysis of the imaging data is hampered by long-tailed distributions and outliers. To combat this, we introduce Wasserstein Geodesic Transformations (WGTs), a new approach that adaptively moves features in Wasserstein space to make downstream analysis less ad-hoc, more stable, and more scalable.

In application to the Cell Painting data, WGTs substantially improve data analysis by enhancing visualization, improving compound clustering, and stabilizing analyses. They also help uncover unwanted spatial effects arising from plate layout, explaining some outlying compound responses. More broadly, the adaptivity of WGT makes it a promising tool a wide-range of cell imaging pipelines.

## 1. Introduction

Cell imaging encompasses a broad class of approaches that use highresolution microscopy to investigate structures of interest in cells and tissues (Lee et al., 2018; Young et al., 2008; Harrison et al., 2023; Scheeder, Heigwer and Boutros, 2018; Zanella, Lorens and Link, 2010). Among other applications, high-throughput cell imaging has been adopted in drug-discovery studies to investigate how candidate compounds change cellular phenotype (Simm et al., 2018; Chandrasekaran et al., 2021; Ziegler, Sievers and Waldmann, 2021). The high-throughput nature means that thousands of compounds can be assessed simultaneously (Rohban et al., 2017; Willis, Nyffeler and Harrill, 2020). The typical pipelines automate the plating, staining, imaging, and feature extraction steps and thereby generate large, diverse sets of morphological features (Caicedo, Singh and Carpenter, 2016; Bougen-Zhukov et al., 2017). These studies can quantify thousands of features covering a host of aspects like morphology, stain intensity, texture, radial distribution, granularity, number of neighbors, and many others (Kamentsky et al., 2011; Singh, Carpenter and Genovesio, 2014).

### 1.1 The Challenge: Long Tails

The imaging features often exhibit problematic properties such as high skewness, long tails, outliers, or high-leverage points (Caicedo et al., 2017). This presents analytical challenges for high-throughput imaging data, in general, and the data we analyze, in particular. This data comprises a Cell Painting study from Wolff et al. (2025) profiling a collection of EU-OPENSCREEN consortium compounds on HepG2 liver cancer cells (c.f. Section 4). Figure 1 plots density estimates for several long-tailed features in this data. The features have been centered and scaled as part of a per-plate -correction approach by subtracting off the median and dividing by the mean absolute deviation from the median. Nonetheless, we still see large measurements in many of the features with values at 20, 30, 50, or even 100 in magnitude. These features also display different types of tails, some features having strong right skewness, left skewness, or being long-tailed yet mostly symmetric. While some of the long tails result from a few outlying observations, often, it is a more fundamental property of the feature’s measurement scale.

**FIG 1.**
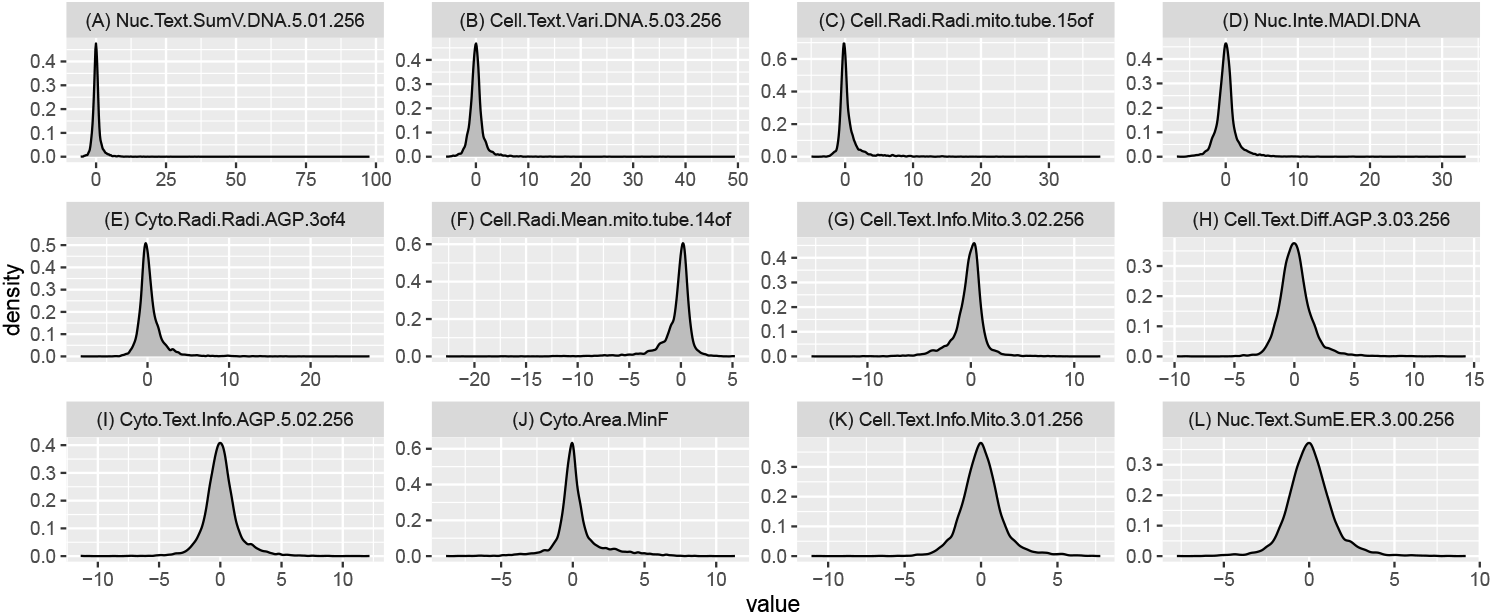
Example density plots of a selection of cell imaging features from Wolff et al. (2025). Names of features have been abbreviated for readability. The subplots are ordered from very long-tailed in (A) to less so in (L).

Poorly behaved features and observations can complicate analysis of high-throughput imaging by distorting visualization or summaries of the data, obscuring the effect of compounds on the cells, and generally undermining the ability to fully process and utilize the panoply of computed features. Consequently, when analyzing high-throughput imaging data, it is often helpful to temper long-tailed features. Issues that arise include, for example,

- Correlation may effectively capture associations among well-behaved features like cell area, but can be misleading when applied to highly skewed intensity or texture features and can be sensitive to a small number of observations.
- Detecting potentially interesting compounds using *z*-scores can be heavily influenced by long tails or outliers.
- Summarizing the relation among compounds using principal component analysis (PCA) can be hampered by skewness or long tails.
- Standard visualizations to explore the biological impact of compounds can become challenging in the presence of outliers or heavily skewed features.

While it’s often wise to temper long-tailed imaging features, a complicating factor is that large values are not always problematic. Sometimes outlying values or features reflect the genuine and strong effect of a particular compound. Consequently, the observations may be informative and warrant closer examination. However, extremely strong effects of one outlying compound can also obscure potentially interesting variation in other other compounds. There’s no free lunch and there is typically a tradeoff between attenuating extreme signals to highlight more subtle ones. However, it is often worthwhile to explore the data under various levels of attenuation to explore the range of signals present. We present an approach to navigate this tradeoff, showing that a transformation which allows tunable attenuation of tails can help illuminate the range of signals in the data from the obvious to more subtle, from highlighting true biology to identifying potentially problematic technical artifacts.

### 1.2. Motivation for a New Approach

Several existing strategies might be used to try and reduce the impact of long tails, for example, winsorizing, censoring, removing outliers, or rank-transforming the data. While scalable, these simple approaches are typically not sufficiently adaptive nor tunable for high-throughput imaging data. Their efficacy varies significantly by feature. For example, censoring the top and bottom 1% may remove noise for some features but exclude many biologically important values for others. Removing high z-score points can have minimal impact on some features but produce substantial missingness in others which can complicate downstream analyses. Censoring or removing points can also distort the data and create artifacts, like bunching of values.

A central issue is the lack of tunability. It is difficult to control how these transformations alter the data. Setting a universal threshold for removing points across all features is rarely appropriate. Similarly, using an aggressive approach to abate skewness like a rank-transformation will often eliminate too much of the original univariate information.

A more adaptive approach is Box-Cox which estimates a power transformation to make the data approximately normal (Box and Cox, 1964; Hunt et al., 2020). While this approach can sometimes work well, Box-Cox has several shortcomings. Not all imaging features can be made approximately normal via power transformations (e.g., mixed discrete/continuous features calculated by thresholding). Box-Cox also often fails to reign in the heavy-tails of symmetric distributions. Most importantly, Box-Cox requires strictly positive values, and extensions to handle non-positive data are often awkward. The method is also highly sensitive to linear rescaling, so results can vary substantially depending on whether preprocessing steps like z-scoring or unit scaling are applied beforehand. This makes Box-Cox particularly sensitive to small changes in analysis pipelines.

These existing transformations struggle with the complexity and variability of highthroughput imaging features. To address this, we propose a more flexible approach guided by four key aims. We discuss these aims conceptually here, and emplore them in more detail in Section 2.

Our first aim (A1) is to attenuate long tails of features in a tunable way:

#### (A1) Tunable Attenuation

Shift feature distributions somewhat toward a more well-behaved target distribution.

In this work, we will use the uniform distribution as our target and thus partially “uniformize” the features. The uniform is a well-behaved distribution with bounded support and thus our partial uniformization will attenuate long, low-density tails and outliers.

While attenuation is paramount, it’s also important to preserve some existing structure in the data and avoid introducing artifacts. Aims 2 and 3 impose two structural constraints:

#### (A2) Monotonicity

The transformation should preserve the ordering of data points.

#### (A3) Density Order Preserving

The transformation should maintain the ordering of densities.

Monotonicity preserves basic ordering structure, which can help interpretation. The Density Order Preserving (DOP) property preserves similar structure for densities. Let *f* be the original density and *f*_*g*_ be the density after a transformation *g*. DOP requires that if *f* (*x*_1_) *> f* (*x*_2_) then *f*_*g*_(*g*(*x*_1_)) *> f*_*g*_(*g*(*x*_2_)) and if *f* (*x*_1_) = *f* (*x*_2_) then *f*_*g*_(*g*(*x*_1_)) = *f*_*g*_(*g*(*x*_2_)).

DOP ensures that relatively dense regions remain relatively dense after transformation, preserving some density related structure like boundedness and number of modes. In particular, DOP helps prevent the introduction of spurious groups that are an artifact of the transformation. For example, transforming a unimodal distribution cannot result in a bimodal one.

For our purposes, these two aims address a common issue: tail-attenuating transformations often introduce density artifacts at the edges of the support. For example, winsorizing piles up values at the minimum or maximum, and strong sigmoidal transformations can compress the tails into a very narrow range, causing density to accumulate near the boundaries (see Supplementary Figure 17). This creates apparent groups (i.e., high-density regions) that were not originally in the data. Aims (A2) and (A3) constrain the transformation to avoid creating these types of spurious grouping artifacts caused by overly aggressive tail suppression. Monotonicity prevents discrete artifacts by ensuring distinct values remain distinct after transformation. Density order preservation goes further, preventing arbitrary density accumulation and thereby requiring that the transformation preserve the presence and number of local and global modes.

Our final aim is to ensure the transformation is robust to linear preprocessing. We realize this as:

#### (A4) Linear-to-Linear Mapping

Applying a linear transformation to the input (e.g., *x → ax* + *b*) should result in (at most) a linear change to the output (e.g., *y → cy* + *d*).

Applying a linear preprocessing step before the transformation should not fundamentally alter the nature of the result. We do not require the output remain exactly the same. For instance, a *z*-score transformation beforehand may shift or scale the transformation result, but any effect on the end result should itself be linear. Fundamentally, we don’t want a linear scale change beforehand to have non-linear effects on the resulting transformed data. Linear changes are mapped to linear changes.

This property is important because linear preprocessing is common in practice (e.g., batch correction or standardization), and we want the transformation to be insensitive to such steps. If a transformation maps linear changes to linear changes, then we can easily make it linearly invariant by centering and scaling its output. We will do this later in our analysis of data so that our results are completely unaffected by any prior linear preprocessing.

Although aims (A1)–(A4) are conceptually simple, none of the commonly used methods satisfy all of them. Table 1 summarizes which properties are met for several common transformations. In the next section we introduce Wasserstein Geodesic Tranformations and show that they achieve all four aims.

**TABLE 1.** Properties of several popular transformation approaches. Checkmark ✓indicates this approach has the property, ∼ indicates the approach partially has the property, and ✗ means the approach lacks the property.

|  | (A1)<br>Tunable Attenuation | (A2)<br>Monotonic | (A3)<br>Dens. Ord. Pres. | (A4)<br>Linear-to-Linear. |
| --- | --- | --- | --- | --- |
| Linear<br>(e.g., $z$-score, 0-1 scaling) | ✗ | ✓ | ✓ | ✓ |
| Censoring<br>(e.g., Winsor, Pt. Removal) | ~ | ✗ | ✗ | ~ |
| Rank Transf | ✗ | ✓ | ~ | ✓ |
| Box-Cox | ✗ | ✓ | ✗ | ✗ |

## 2. Wasserstein Geodesic Transformations

Let *X* denote one of our univariate image features and let *F* be its CDF. Let *F* ^*∗*^ be the CDF of some well-behaved target distribution. (Ultimately, this will be a uniform distribution, but for now we keep it generic.) We would like to move *F* somewhat towards *F* ^*∗*^ to partially attenuate *F*. Define the transformation *g*_*θ*_(*x*) as

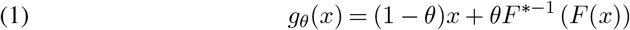

for some *θ* ∈[0, 1]. We will show that transformations of this form are very natural choice if we wish to achieve (A1)–(A4).

The form of *g*_*θ*_ can be motivated by considering how to develop a family of transformations that allow broad tuning of (A1). To see this, let *F*_*g*_ be the CDF of *X* under some transformation *g*. Aim (A1) seeks to move *F*_*g*_ somewhat towards *F* ^*∗*^. A reasonable approach for doing this is to define some family of potential transformations 𝒢 and search over *g* ∈ 𝒢 to find a transformation that moves *F*_*g*_ away from *F* and towards *F* ^*∗*^the desired amount. A capacious way to do this is to let 𝒢 parameterize a curve {ℱ = *F*_*g*_ for *g* ∈𝒢 } between *F* and *F* ^*∗*^. Given such a 𝒢 we can then tune *g* to place *F*_*g*_ at any point along the curve between *F* and *F* ^*∗*^.

There are many ways to build such a curve. In this work we form ℱ by analogy to connecting two real vectors *x* and *y* with the path carved by their *θ*-weighted average over 0 ≤ *θ* ≤ 1. We generalize this idea and build a curve ℱ between *F* and *F* ^*∗*^ by defining some metric *d* on our space of distributions and let *F*_*θ*_ be the *θ*-weighted Fréchet mean of *F* and *F* ^*∗*^ according to *d*:

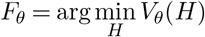

so that *F*_*θ*_ minimizes the weighted Fréchet variance *V*_*θ*_(*H*) = (1 −*θ*)*d*^2^(*H, F*) + *θd*^2^(*H, F* ^*∗*^). Given a choice of *d* this will parameterize a curve ℱ = {*F*_*θ*_, 0 ≤ *θ* ≤ 1} such that *F*_0_ = *F* (corresp. to *g*_0_(*x*) = *x*) and *F*_1_ = *F* ^*∗*^ (corresp. to *g*_1_(*x*) = *F* ^*∗*−1^(*F* (*x*))).

Different curves will be constructed for different metrics *d*. For example, if *d* is the *L*^2^ distance 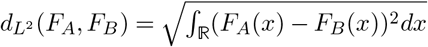 then *F*_*θ*_ is the mixture of *F* and *F* ^*∗*^ so that 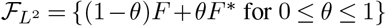 for 0 ≤ *θ* ≤ 1}. These mixture distributions are not what we seek as they entail mixing CDFs of features with potentially different supports. A better distance to use is the Wasserstein 2-distance

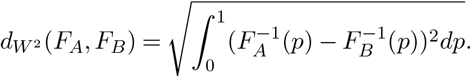

In this case the Fréchet mean will form distributions where the quantile functions (inverse-CDFs) of *F* and *F* ^*∗*^ are mixed:

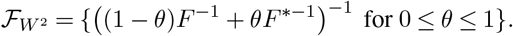

This mixing makes sense as the quantile functions have the same domain of [0, 1].

This curve 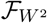 is the geodesic path between *F* and *F* ^*∗*^ under this Wasserstein metric (Santambrogio, 2015; Panaretos and Zemel, 2020). This Wasserstein geodesic corresponds to the parameterized family of transformations:

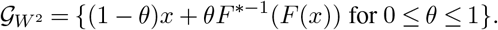

This is precisely the proposed family of transforms in Equation 1. While one can define such a family for any target distribution *F* ^*∗*^, in this work we consider a *U* (0, 1) target. This will move the data distribution *F* somewhat towards a well-behaved uniform distribution. Consequently, we will work with the family

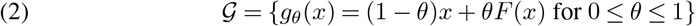

which is precisely the Wasserstein geodesic between *F* and a *U* (0, 1) target distribution *F* ^*∗*^. Consequently, we will call such transformations Wasserstein Geodesic Transformations (WGTs).

### 2.1. Properties

We will now explore how WGTs following Equation 2 achieve aims (A1)–(A4).

#### 2.1.1. A1: Tunable Attenuation

Aim (A1) seeks to move *F*_*θ*_ somewhat closer to a well-behaved target. In particular, we want to partially uniformize the distribution by moving it towards that of a *U* (0, 1). One can show that if *F*_*U*_ is a standard uniform distribution then 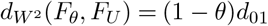 and 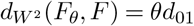 where 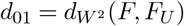. (Details may be found in Supplementary Section 6.1.) Thus as *θ* travels from 0 to 1 we have that *F*_*θ*_ gets closer to a *U* (0, 1) distribution and further away from the original distribution.

#### 2.1.2. A2: Monotonicity

The proposed family of transformations preserves ordering. This follows since for any *θ <* 1 the transformation *g*_*θ*_ is increasing since *x* is increasing and *F* is non-decreasing. When *θ* = 1 we have *g*_*θ*_ = *F* which is always non-decreasing and sometimes increasing. Thus *g*_*θ*_ is strictly increasing for *θ <* 1 and at least weakly increasing for *θ* = 1.

#### 2.1.3. A3: Density Order Preserving

Among the many possible targets *F* ^*∗*^ to which we could transform a feature’s distribution, the uniform distribution has a unique advantage of having the DOP property. Without loss of generaity we choose *F* ^*∗*^ to be *U* (0, 1). Another way of phrasing the DOP property is that that *f*_*g*_(*g*(*x*)) is a strictly increasing function of *f* (*x*). This property helps prevent the transformation from introducing spurious groupings due to excessive tail suppression. Not only does a uniform target guarantee DOP, it is the only target for which WGTs universally preserve density order across arbitrary features (see Supplementary Section 6.2).

Beyond this, the uniform distribution is particularly well-behaved as a transformation target. It means the WGTs somewhat spread density more evenly across the support, suppressing long, sparse tails and outliers. This “partial uniformization” mitigates outlying feature values and enhances underlying signal that might otherwise be obscured. We examine this effect empirically in Section 4, and show analytically in Supplementary Section 6.3 that WGTs reduce tail heaviness across a broad class of distributions.

#### 2.1.4. A4: Linear-to-Linear Mapping

WGTs are essentially insensitive to linear preprocessing of the features. For any *x* let *y* = *ax* + *b* for some *a >* 0 and let *F*_*X*_ and *F*_*Y*_ be the corresponding CDFs. If

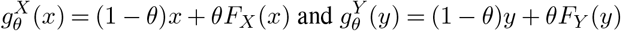

are the corresponding WGTs, we can show that

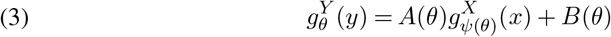

where *ψ*(*θ*) = *θ/*(*a*(1 −*θ*) + *θ*), *A*(*θ*) = *a*(1 −*θ*) + *θ*, and *B*(*θ*) = *b*(1 −*θ*). (See Supplementary Section 6.4.)

Equation (3) shows that the *Y* -geodesic transformation is equivalent to (i) considering an *X*-geodesic transformation but instead of using *θ* we use *ψ*(*θ*), and then (ii) linearly re-scaling the resulting output using *A*(*θ*) and *B*(*θ*). After reparameterizing, a linear scale change beforehand becomes a linear scale change afterwards. Moreover, Section 3.2 will show that we can remove the reparameterization step by choosing *θ* in a linearly-invariant manner.

The capacity of the transformation family is essentially unaffected by linear preprocessing. This is a strong and distinctive property. Most other transformation methods do not share this. For example, Box-Cox results will depend on whether the data was scaled using the standard deviation or IQR. Such choices do not affect WGTs except for changing a final linear scale. Typically, we will *z*-score features after transformation and thus linear preprocessing makes zero difference for WGTs.

## 3. Fitting and Tuning

We apply WGTs to high-throughput imaging data in an unsupervised context. Given *N* feature measurements *X* = (*x*_1_, …, *x*_*N*_), we must estimate 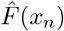 and select a good *θ*^*∗*^. The transformed values are then

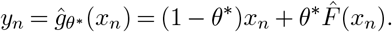

### 3.1. Estimating F

For continuous measurements we can let 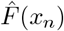 be the empirical CDF (eCDF) so that

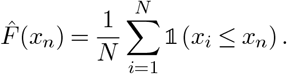

However, this estimate may have undesirable properties for discrete or mixed discrete-continuous features, for example, those created via thresholding. For features with unique values, the eCDF yields an empirical uniform distribution with spacing 1*/N*. For discrete or mixed-type features, however, the eCDF introduces gaps proportional to value repetition. In Figure 2, we show *N* = 1000 samples from *X* = (*X*_0_ − mean(*X*_0_))*/*sd(*X*_0_) with *X*_0_ = log(1 + *Z*) and 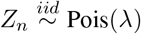. Subplot (A) shows that when *λ* = 10, the eCDF-transformed data (red) is not uniform and has visible jumps reflecting repeated values. In subplot (B), *λ* = 10^6^ results in nearly all unique values, and so the eCDF output is approximately uniform. Figure 3 shows a different case: an even mixture of *N* (0, 1) and a point mass at 0. The resulting eCDF-transformed feature exhibits a large gap at 0 which is likely undesirable as it is essentially an artifact of the transformation.

**FIG 2.**
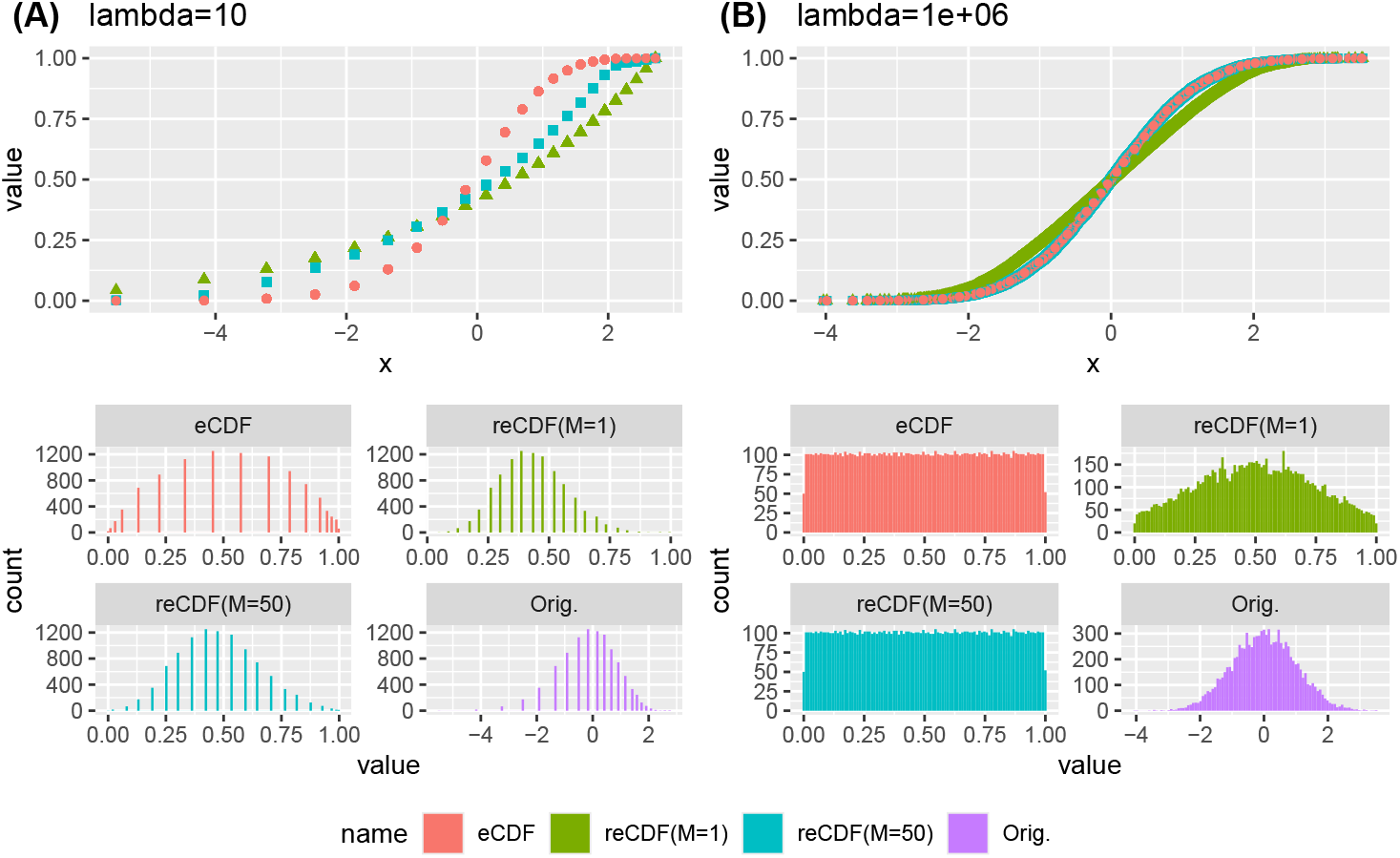
Examples of eCDF and reCDF estimates of 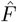 for a log-transformed Poisson distribution. Subplot (A) consider a case where λ = 10 and subplot (B) considers a case where λ = 106. The top of each figure displays the CDFs evaluated at each point. The bottom displays histograms of the transformed values.

**FIG 3.**
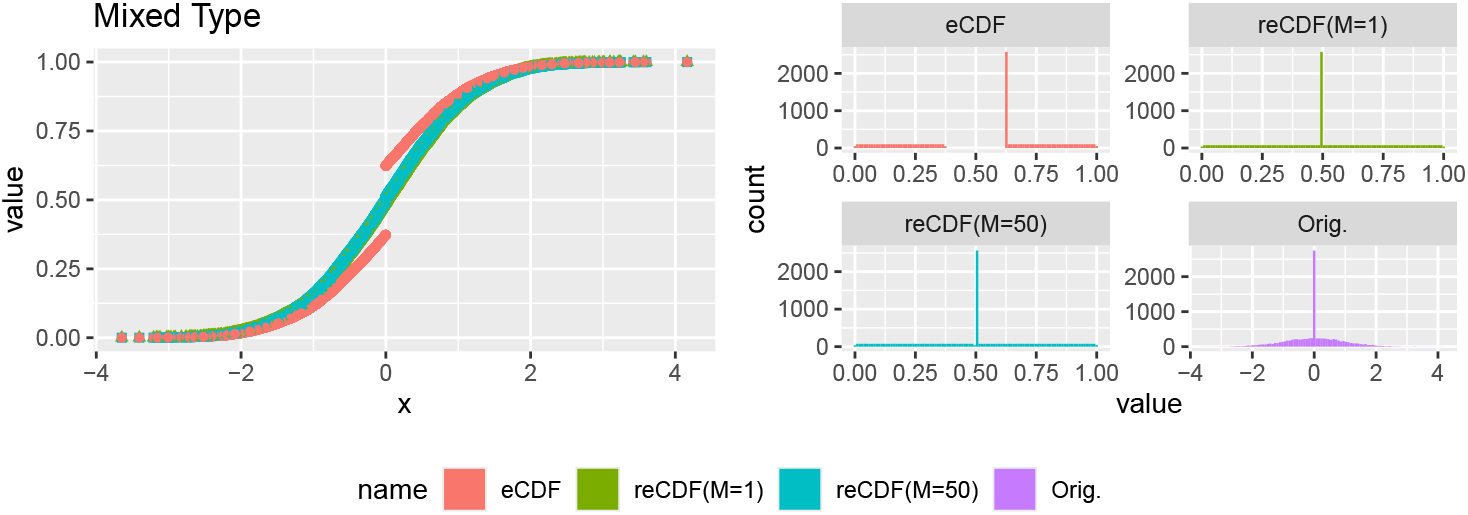
Examples of eCDF and reCDF estimates of 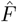 for a mixture of a Normal and degenerate distribution. Left displays the CDFs evaluated at the support points. Right displays histograms.

To address this issue, we estimate 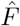 using a regularized empirical CDF (reCDF). Let *W* = *{w*_1_, …, *w*_*Q*_*}* be the set of *Q* ≤ *N* unique values among *x*_1_, …, *x*_*N*_ and 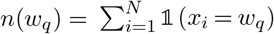 be the number of *x*_*i*_ exactly equal to *w*_*q*_ ∈ *W*. For a regularization parameter *M* ≥ 1 define the regularized cumulative count at *x*_*n*_ as

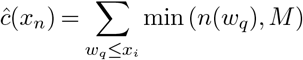

and the regularized maximum count as

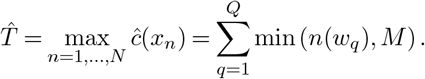

Then the regularized CDF estimate 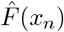 is

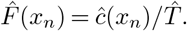

This regularized CDF caps the jump size at any point to *M*. It matches the standard eCDF when all values are unique or when *M* ≥ max_*q*_ *n*(*w*_*q*_). For non-distinct values, smaller *M* regularizes the estimate. For example, *M* = 1 yields the proportion of unique values less than or equal to *x*_*n*_. Thus, *M* controls the maximum jump in 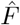 and the size of gaps in the transformed feature’s support. In our later analysis of real data we use the reCDF estimate using *M* = 1.

Revisiting Figure 2 and Figure 3 one can consider the reCDF transformed values using *M* = 1 (green) and *M* = 50 (blue). For small *M*, the reCDF can differ markedly from the eCDF, though both flatten in the tails. As repeated values decrease, reCDF and eCDF estimates converge. While *M* = 1 does not yield a uniform distribution it still reigns in the tails and evenly spaces the support. Notably, in Figure 3 neither versions of the reCDF create a grouping artifact like the eCDF.

### 3.2. Determining θ

We want the transformation to produce essentially the same result whether applied to *X* or *Y* = *aX* + *b*. Any difference should be limited to a linear scaling, which can be corrected afterward (e.g., using a *z*-score). This affects how we choose *θ*, since the meaning of *θ* depends on the scale of the input variable. Previously, we saw that if *X*_*θ*_ = (1 − *θ*)*X* + *θF*_*X*_(*X*), and *Y*_*θ*_ = (1 − *θ*)*Y* + *θF*_*Y*_ (*X*) then

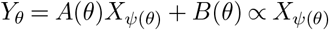

where *ψ*(*θ*) = *θ/*(*a*(1 − *θ*) + *θ*). That is, scaling *X* re-parameterizes the transformation according to *ψ*(*θ*). Thus *X*_*θ*_ and *Y*_*θ*_ need not be perfectly correlated; but *Y*_*θ*_ and *Xψ*(*θ*) will be. Consequently, if we want the *X* and *Y* transformations to be linear re-scalings of each other then if 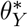 is the chosen value for *θ* using *Y* then 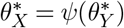 should be the parameter value chosen using *X*.

One way to ensure this is to let *θ*^*∗*^ be the value of *θ* such that

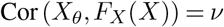

for some chosen correlation *ν*. (This is unique since it is strictly increasing in *θ*, c.f. Supplementary Section 6.5.) This procedure is scale invariant since if 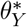 is the chosen value using *Y* then and since *F*_*X*_(*X*) = *F*_*Y*_ (*Y*) and 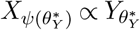 then

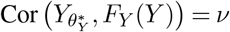

and thus by uniqueness 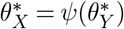.

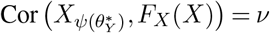

This approach has a closed-form solution for *θ*^*∗*^:

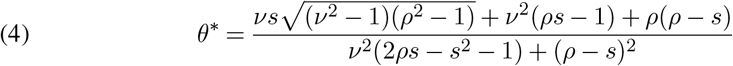

where *ρ* = Cor(*X, F* (*X*)) and *s* = Var(*F* (*X*))*/*Var(*X*). (See Supplementary Section 6.5.) Note that the correlation parameter *ν* must be between *ρ* and 1 which will change from one feature *X* to another. Thus, instead of working directly with *ν*, we re-parameterize with a parameter *η* ∈ [0, 1] letting

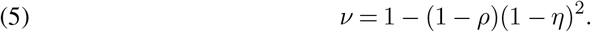

This *η*-parameterization has a nice interpretation. Let Δ be a linearly-invariant version of the Wasserstein distance defined for two distributions *F*_*A*_ and *F*_*B*_ so that

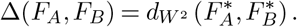

where 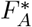 is the CDF of 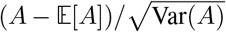 where *A* ∼ *F*_*A*_, and similary for 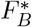. Further define the normalized (linearly-invariant) Wasserstein distance between the distribution *F*_*θ*_ of *X*_*θ*_ and the uniform distribution *F*_*U*_ of *F* (*X*) as

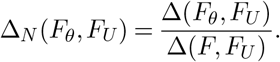

Then 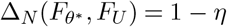 (see Supplementary Section 6.5), so *η* directly controls a normalized, linearly-invariant Wasserstein distance between *X*_*θ*_ and the uniform target. Setting *η* near zero keeps *X*_*θ*_ close to *X*, while setting *η* near one brings it closer to a uniform. The resulting distribution (up to linear scaling) is invariant to linear preprocessing of *X*.

In practice, *η* is user-chosen. We estimate *s* and *ρ* as 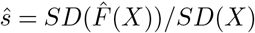 and 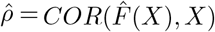 using standard sample estimates, then compute *θ*^*∗*^ by substituting these into Equations 4 and 5. For downstream analyses one may experiment with different values of *η* ∈[0, 1] to tune different levels of feature attenuation, or pick a fixed value like *η* = 3*/*4 for all features.

## 4 Results

We are interested in improving the analysis of a recent high-throughput imaging dataset from Wolff et al. (2025), which profiles a large number of compounds from the EU-OPENSCREEN consortium. Aimed at supporting small molecule research, the study emphasizes assay quality and provides a comprehensive, high-quality screening resource.

The authors used 384-well plates to profile HepG2 human liver cancer cells with the Cell Painting assay. We focus on sixteen plates from the study, which screened 355 compounds from the EU-OPENSCREEN Bioactive set. Each experiment was run in quadruplicate across four sites in Czechia, Germany, and Spain, yielding sixteen replicates per compound. DMSO wells served as negative controls, while tetrandrine and nocodazole were used as positive controls. Approximately 2800 image features were extracted using CellProfiler, based on six fluorescent stains targeting eight cellular regions: DNA, cytoplasmic RNA, nucleoli, actin, Golgi apparatus, plasma membrane, endoplasmic reticulum, and mitochondria. A few degenerate features were excluded, usually due to CellProfiler processing issues like overly-aggressive image thresholding.

While high-quality, this data can still benefit from the robustifying effects of WGT. In our analysis we apply a series of transformations (including WGT) on a plate-by-plate basis. To reduce plate-to-plate batch effects, we normalize each plate by subtracting the median and dividing by the mean absolute deviation from the median.

### 4.1. Attenuation and Visualization

As noted in Section 1.1, a common challenge in analyzing imaging data is the presence of long-tailed features or outlying observations. While reducing long tails can often improve analysis, the dilemma is that large observations are not always problematic; in some cases, outlying values or features may reflect genuine biological variation of interest. In Figure 4, we show boxplots of four features with outlying observations, stratified by compound along the *x*-axis. Subplots (A) and (B) illustrate features with minimal biological signal and clear outliers. In these cases, the outlying values are unlikely to reflect biology, so attenuating them may help uncover underlying signal of interest. In contrast, subplots (C) and (D) show features with strong biological signal in some compounds, weaker signal in others, and a few clear non-biological outliers. For such features, there can be a tradeoff: largely, we want to preserve strong biological patterns while still somewhat dampening outlying values that may obscure weaker signal or reflect noise. Our results will demonstrate that this tradeoff can often be navigated advantageously using WGT. A slight reduction of the most obvious signal is often a worthwhile price for helping uncover otherwise obscured biology.

**FIG 4.**
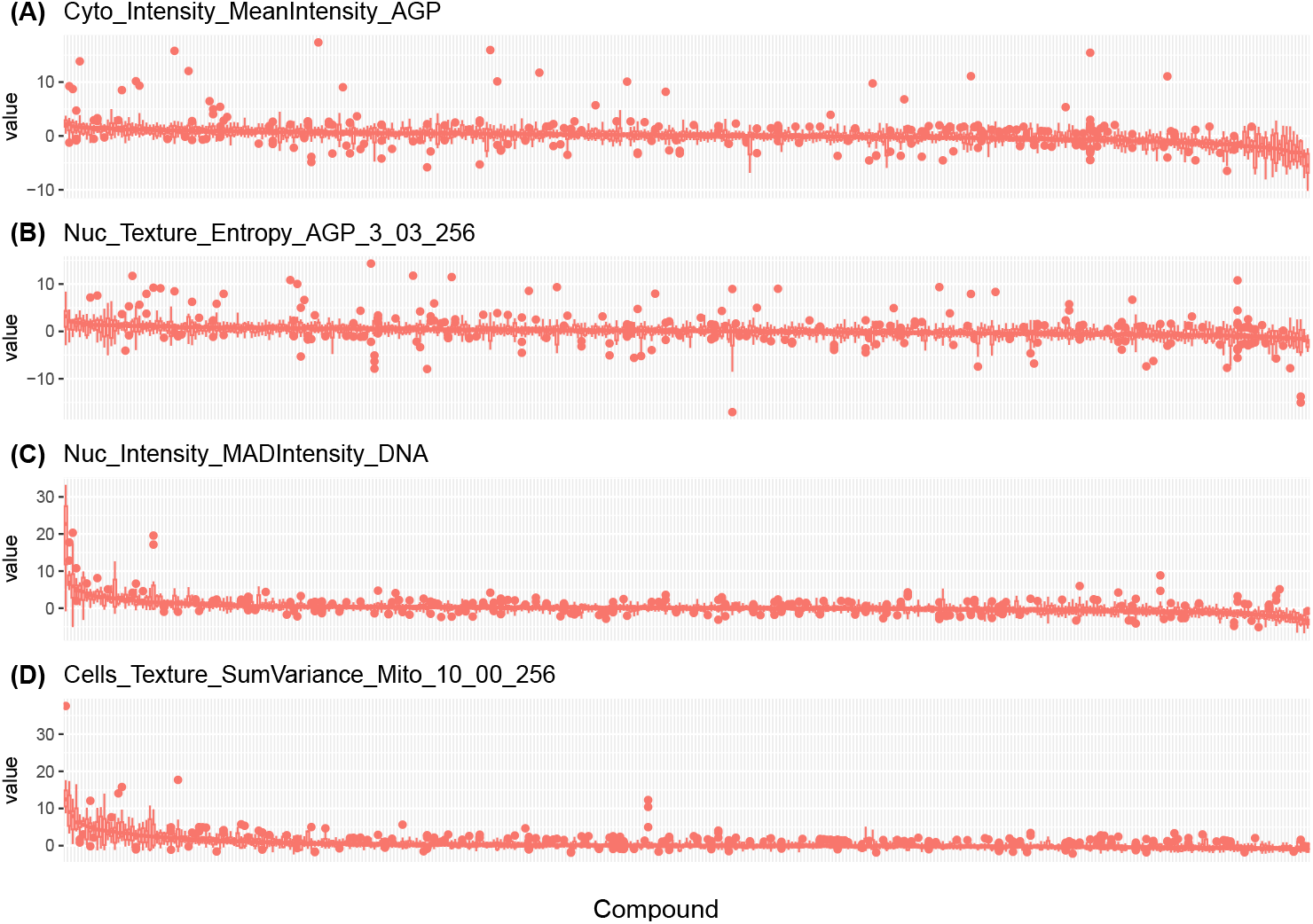
Boxplots of four long-tailed features across compounds. The x-axis is ordered decreasing according to median and is different for each subplot. Subplots (A) and (B) show weak signal with outliers. Subplots (C) and show stronger biological signal with some non-biological outliers.

To help navigate this tradeoff WGT allows tuning attenuation strength via *η*. In general, one may experiment with several values of this parameter throughout the course of an analysis. Empirically, we find that small changes in *η* do not change the results much (e.g., *η* = 0.70 v. 0.75); however the results will substantially change over a wider range (e.g. *η* = 0.1 v. *η* = 0.9). To illustrate how *η* tunes strength, Figure 5 shows density plots of WGT-adjusted values for two long-tailed features over a range of values of *η*. As *η* increases, the transformation progressively suppresses the tails of the distribution, concentrating more on the bulk of the data. Thanks to the density order preserving property, some of the features of the distributions’ shapes are preserved: right-skewness remains visible (though reduced) in subplot (A), while the approximate symmetry is maintained in subplot (B). The choice of *η* controls a tradeoff. Larger *η* attenuate tails to emphasize more typical values, while smaller *η* highlights tails more.

**FIG 5.**
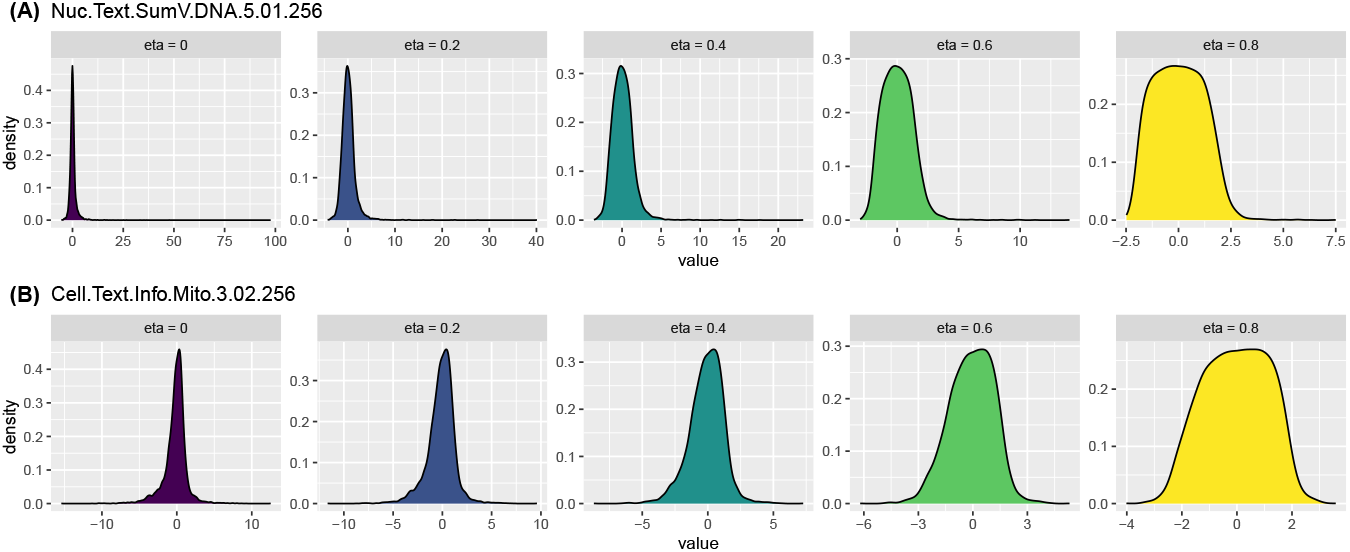
Density plots of WGTs for two long-tailed features over a range of values of η.

There is no universally optimal choice of *η*, and we encourage trying different values to explore how the transformation emphasizes various signals in the data. However, as shown in Figure 5, for this analysis we find that the long tails of some distributions are not sufficiently reduced until *η* ≳ 0.5. To avoid repeating similar analyses across many values of *η*, we use *η* = 0.75 for the remainder of this work as a practical compromise. This value generally strikes a good balance between attenuating outliers and preserving signal, though it should not be treated as a fixed rule.

A quick way to assess the effect of our chosen attenuating transformations is to compute the percentage of outliers for each feature. A consequence of tail attenuation is a substantial reduction in the number of outliers across features. In Figure 6, we compute the percentage of values identified as outliers (more than 1.5 IQR above or below the first or third quartile) for all *∼* 2800 features. To give a point of comparison with a more traditional transformation, we also apply a Box-Cox transformation to all features, using a standard modification of Box-Cox to accommodate negative values following Bickel and Doksum (1981) and replacing exact zeros with 1 *×* 10^−10^. Box plots in Figure 6 summarize the percentages of outliers across features for each transformation method. Without any transformation, this rule typically flags many observations as outliers, with a median around 5-10% of points and about 75% of features having less than 15% outliers. The Box-Cox transformation provides a modest reduction in identified outliers. In contrast, WGT dramatically lowers the percentage, with a median of 0.25% and third quartile of 1%. This reduction not only helps limit the influence of outliers, but also makes it easier to analyze data without as much need to filter, adjust, or worry about their disproportionate impact. If we do remove outlying points for subsequent analyses, WGT allows us to remove many fewer, making it easier to identify and remove the truly extreme points that would otherwise cause issues for analyses. Nonetheless, a key advantage of the WGT transformation is that it attenuates the influence of outlying values without requiring their outright removal, which is helpful when processing thousands of very disparate features while ensuring that we don’t inadvertently lose important information.

**FIG 6.**
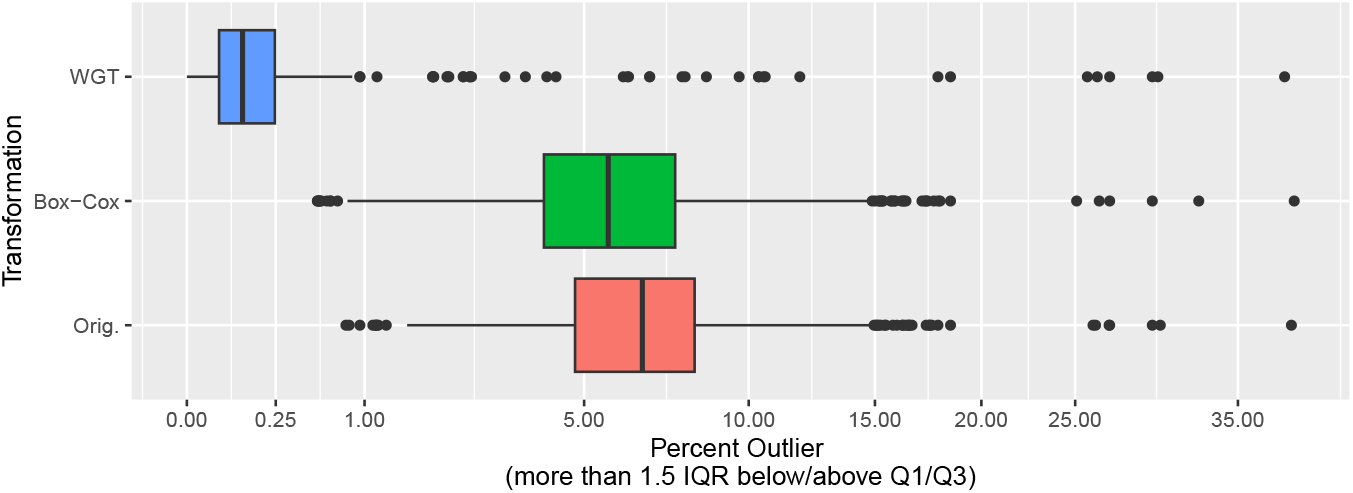
Percentage of outliers for no transformation, Box-Cox, and WGT(η = 3/4). Each data point contributing to the box-plots represents the percentage of outliers for one of the ∼ 2800 variables.

Another good sanity-check for the transformations is to make some density plots of features. Figure 7 does this, comparing the resulting feature distributions across three approaches: no transformation, Box-Cox, and WGT with *η* = 3*/*4. The comparison in Figure 7 highlights key differences in flexibility and tunability. While Box-Cox reigns in the tails, its effect is generally weaker than that of WGT and often very small for symmetric distributions, as seen in subplots (C) and (D).

**FIG 7.**
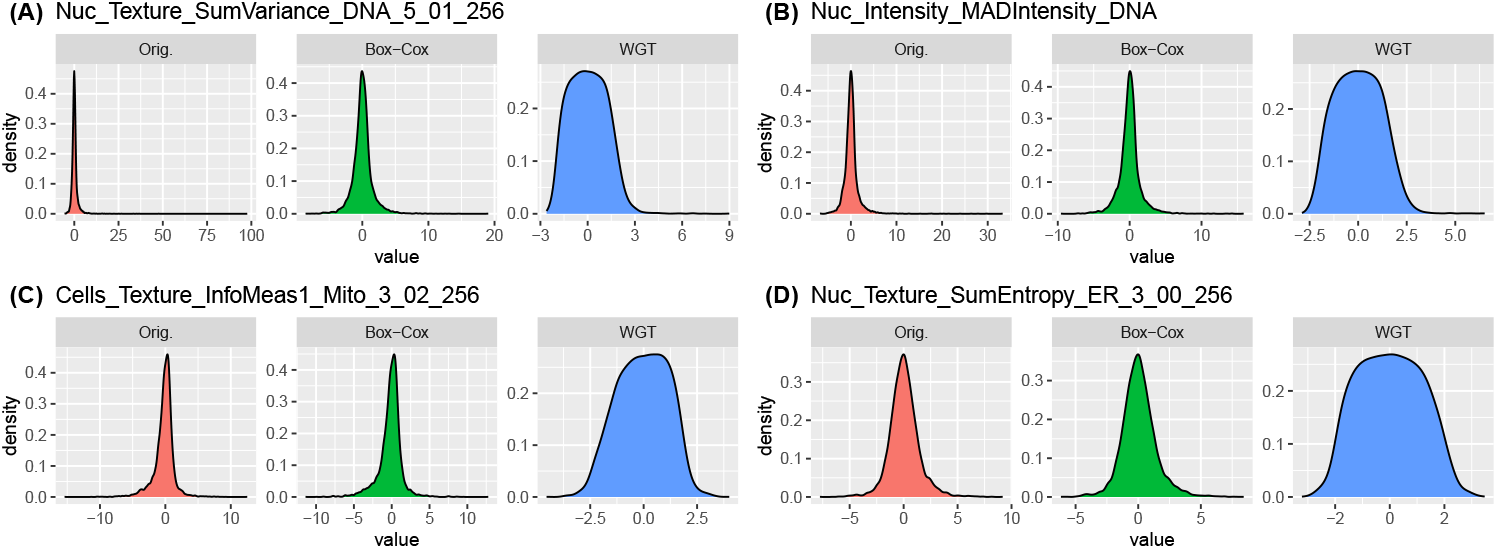
Comparison of univariate densities for no transformation, Box-Cox, and WGT(η = 3/4) for several features.

An immediate benefit of WGT is that its attenuation of long tails can enhance visualization. As an example analysis, in Figure 8 we show boxplots of several imaging features comparing values for EU-OPENSCREEN (EOS) compounds against negative controls (DMSO) and positive controls (nocodazole and tetrandrine). WGT makes visualization more straightforward. For instance, in subplot (A), the feature Nuc_Texture_SumVariance_DNA_5_01_2 56 has a long-tailed distribution in the EOS group, making it difficult to discern group differences. A Box-Cox transformation improves this slightly, but WGT more clearly reveals that nocodazole produces substantially lower values, with a similar (though less pronounced) effect for DMSO. This clarity in group separation is not specific to one feature; subplots (B)-(D) illustrate that WGT generally improves the ease of these types of visual comparisons. This highlights an advantage of WGTs: we can produce more informative visualizations without removing any data or resorting to bespoke adjustments for problematic features. This is particularly useful for analysis of the thousands of high-throughput imaging features at scale.

**FIG 8.**
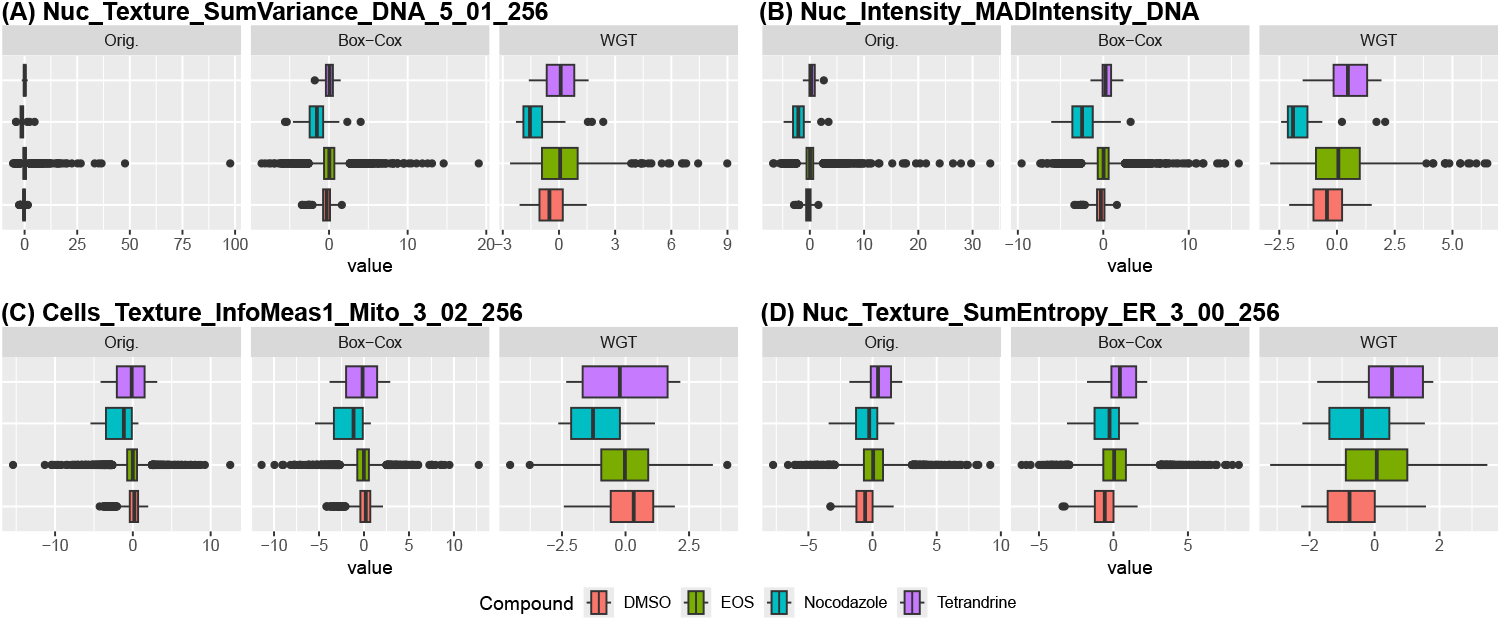
Boxplots of four imaging features comparing values for EU-OPENSCREEN (EOS), negative controls (DMSO), and positive controls (nocodazole and tetrandrine).

Building on the compound-stratified analysis in Figure 4, Figure 9 shows the same features after applying WGT. Subplots (A) and (B) are notably more interpretable post-transformation; attenuating non-biological outliers reveals clearer compound-to-compound differences. A similar improvement is seen in subplots (C) and (D). While the most highly expressed compounds are somewhat less large after WGT, they remain distinct and clearly biologically meaningful. Crucially, they no longer dominate the scale of the plots, allowing subtler signals in other compounds to emerge visually and, as we will show later, quantitatively. The outliers now flagged in the boxplots are less large, less visually disruptive, and seemingly less likely to represent true biological variation. (A similar plot for Box-Cox may be found in Supplementary Figure 18, which shows some improvement, but not as much as WGT.)

**FIG 9.**
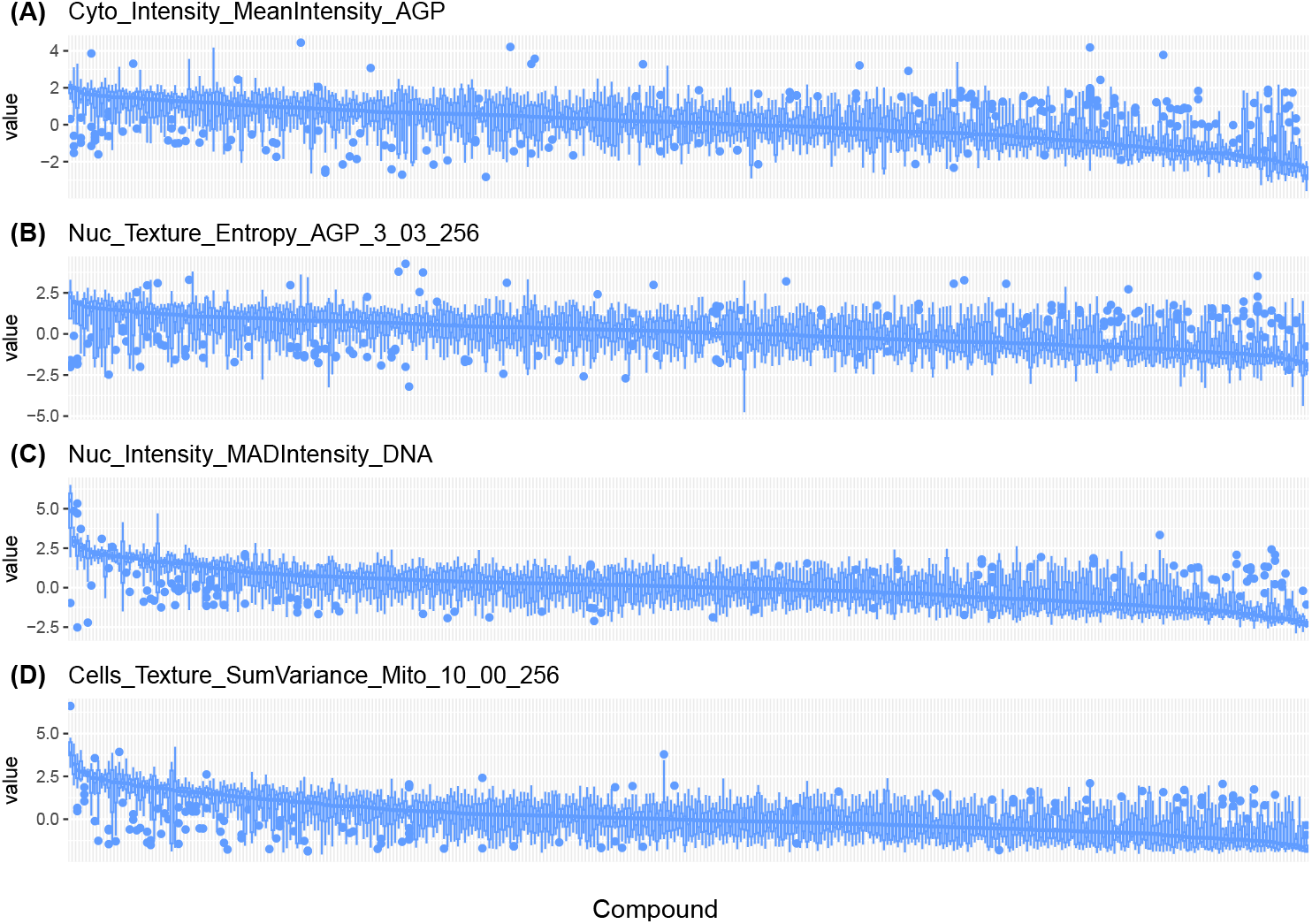
Re-analysis of features in Figure 4 after WGT. The x-axis is ordered decreasing according to median and is different for each subplot.

### 4.2. Correlation and Sensitivity

A major benefit of attenuating transformations is their ability to stabilize analyses by reducing sensitivity to a small number of outliers or features. Figure 10 illustrates this point by presenting 2D histograms of several feature pairs. These plots demonstrate that data transformation can significantly affect common summary metrics such as correlation.

**FIG 10.**
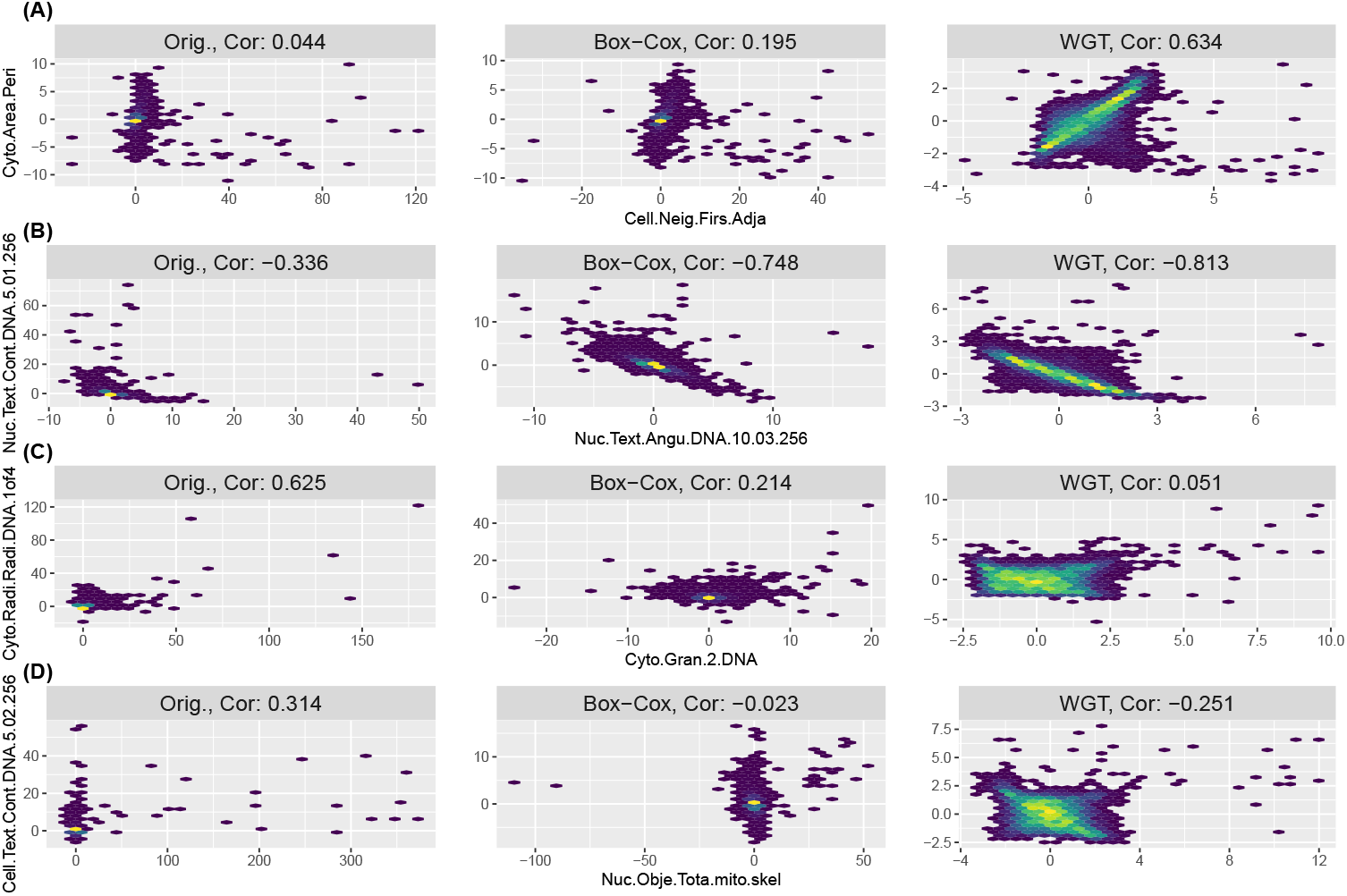
Effect of data transformation on pairwise correlations between example imaging features. WGT attenuates long tails, revealing or reducing linear relationships that are otherwise obscured or exaggerated in the raw data.

In subplot (A), we examine the relationship between Cells_Neighbors_FirstClosestDista nce_Adjacent and Cyto_AreaShape_Perimeter. Without transformation (left column), these features show near-zero correlation, driven largely by the long tail of the perimeter feature. Applying a Box-Cox transformation moderates the outliers, increasing the correlation to approximately 0.19. WGT more effectively suppresses the tail, yielding a stronger correlation of about 0.63. WGT identifies that (excluding the tail) there is a meaningful linear relationship between these features. This is more difficult to identify in the unadjusted data.

Subplot (B) presents similar story comparing Nuc_Texture_AngularSecondMoment_D NA_10_03_256 vs. Nuc_Texture_Contrast_DNA_5_01_256. The untransformed features have a weak negative correlation of − 0.336, which strengthens to −0.74 after a Box-Cox transformation and to −0.81 with WGT. WGT identifies a strong negative relationship between the bulk of these two features by reducing the influence of high-leverage values.

While sometimes a tail-attenuating transformation can strengthen the linear relationship, we should not always expect this to be the case. Subplot (C) shows a case where transformation reduces apparent association. Here, we compare Cyto_Granularity_2_DNA and Cyto_ RadialDistribution_RadialCV_DNA_1of4. The untransformed correlation is 0.62, but after attenuating the tails, Box-Cox reduces this to 0.21 and WGT further to 0.05, indicating the strong correlation was primarily driven by the tails of the distributions.

Finally, subplot (D) illustrates that tail attenuation can even reverse the direction of correlation. The pair Nuc_ObjectSkeleton_TotalObjectSkeletonLength_mito_skel vs. Cells_T exture_Contrast_DNA_5_02_256 shows a weak positive correlation (0.31) before transformation, but WGT shifts it to a weak negative correlation (− 0.25). Box-Cox is in between (correlation near zero).

Overall, Figure 10 underscores how data transformations can substantially impact common metrics like correlation. While it may be important to investigate outlying measurements, many standard metrics are highly sensitive to such values. WGTs help clarify whether a pattern is representative of the bulk of the distribution or an artifact of a small number of values. This is especially critical in high-throughput imaging data, where thousands of features must be analyzed without manual inspection. Although robustified metrics exist, transformations like WGT offer a more general way to enable the application of traditional statistical tools without undue influence from long-tailed measurement scales or outlying observations.

To assess the sensitivity of correlation estimates, Figure 11 examines how correlations behave under repeated sub-sampling. For four representative feature pairs, we compare results under no transformation, a Box-Cox transformation, and a WGT with *η* = 3*/*4. In each case, we randomly sub-sample 10% of the data in 100 Monte Carlo replicates and compute the resulting correlations. The distributions of these estimates are summarized using boxplots.

**FIG 11.**
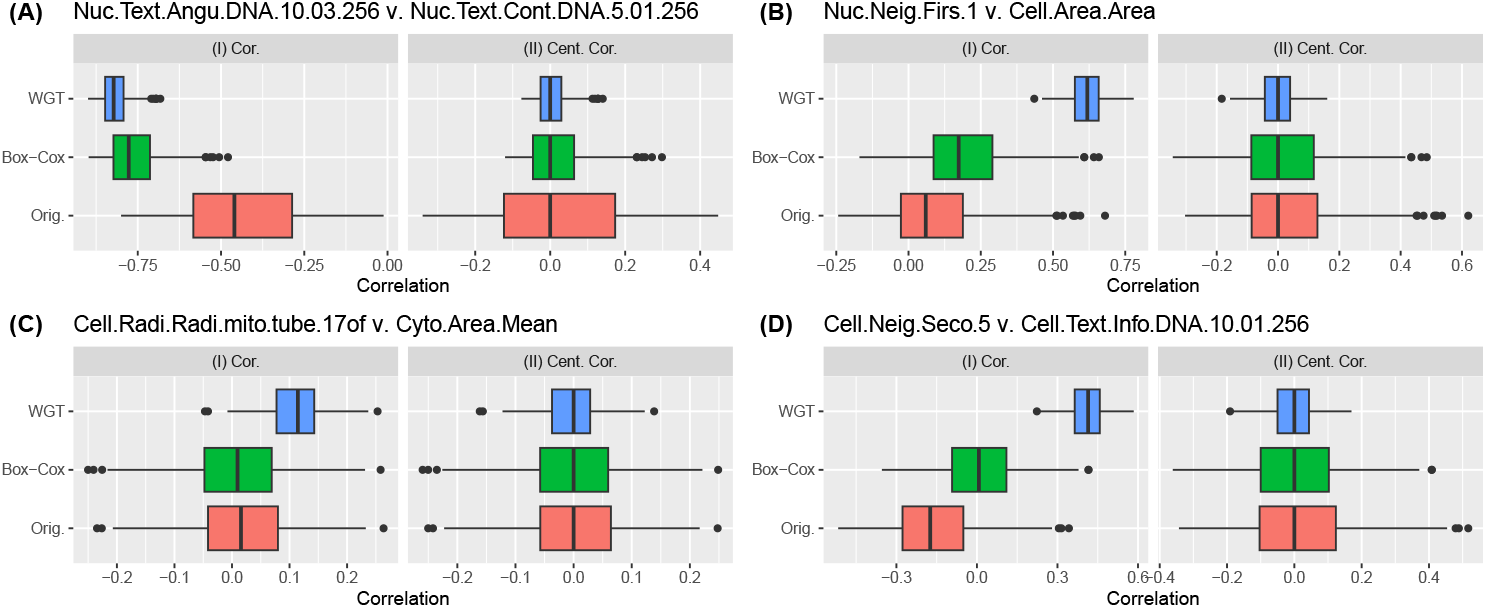
Stability of correlation estimates under sub-sampling across four feature pairs. WGT consistently reduces variability and improves robustness compared to no transformation or Box-Cox, highlighting its utility in attenuating the influence of outlying values.

Subplot (A) shows the correlation between Nuc_Texture_AngularSecondMoment_DNA _10_03_256 and Nuc_Texture_Contrast_DNA_5_01_256. Looking at panel (I), the WGT transformation yields stronger and more stable correlation estimates compared to the untransformed or Box-Cox-transformed versions. To highlight differences in variability, panel (II) shows the centered distributions (after subtracting the median), making it clear that WGT sharply reduces the spread of correlation estimates. While Box-Cox slightly reduces variation, the effect is not nearly as strong or consistent as WGT. Subplots (B)-(D) demonstrate that this increased stability is consistent across diverse feature pairs. In general, we can see that tail-attenuating transformations like WGT will substantially increase the stability of metrics like correlation.

### 4.3. Dimensionality Reduction and Clustering

WGT also proves valuable for exploratory analyses such as dimensionality reduction, which benefits from the same strengths we’ve previously observed: improved visualization and more meaningful inter-feature correlations. Figure 12 illustrates this by showing the first two principal components (PCs) for three different preprocessing approaches: no transformation, Box-Cox, and WGT. To assess the robustness of these visualizations, we repeatedly subsample the dataset, each time retaining a random selection of 10 compounds and 10 image features, and then compute the PCs. Subplots (A)–(C) show three representative examples from these resampled datasets.

**FIG 12.**
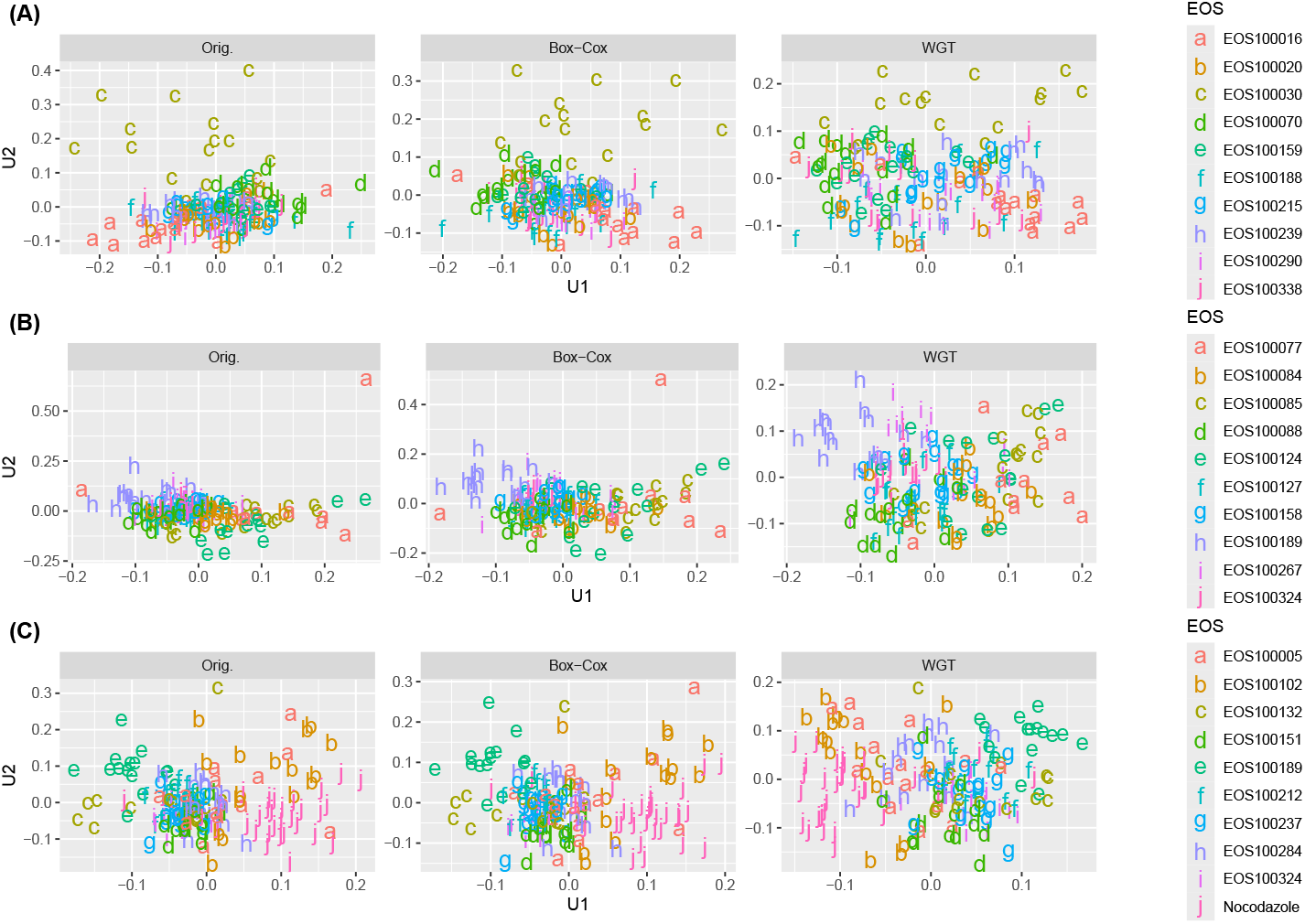
Principal component plots from three representative subsamples illustrate the effect of preprocessing on compound separation. Compared to no transformation or Box-Cox, WGT more consistently spreads out the data, reduces the influence of outlying points, and enhances visual separation of compound clusters.

In Subplot (A), the untransformed and Box-Cox data both compress the points into a small region, primarily reflecting variation from a single compound. In contrast, WGT distributes the data more evenly across the plot, allowing clearer visual separation among different compounds. Subplot (B) shows a case where the untransformed data are heavily influenced by the presence of one or two outlying points, which dominate the PCs and obscure overall structure. While Box-Cox mitigates this effect to some extent, WGT more effectively downweights these points, revealing a clearer clustering pattern among the compounds. Finally, in Subplot (C), both the untransformed and Box-Cox data produce reasonably spaced clusters. However, WGT further enhances this.

To quantitatively evaluate clustering performance, Figure 13 presents several clustering metrics computed from principal component analyses on 100 Monte Carlo subsamples of the data, as described previously. For each subsample, we calculate the first *K* principal components with *K* = 1, 2, 3, 4, 5, and 10, then assess clustering using four metrics: CalinskiHarabasz (CH), Dunn, Silhouette, and the *tr*(*W* ^−1^*B*) metric (Desgraupes, 2023). The bold lines show the median values across subsamples, while the shaded ribbons indicate the 25^*th*^ and 75^*th*^ percentiles. Generally, clustering metrics quantitatively balance within-cluster compactness against between-cluster separation. WGT tends to spread points more evenly across the space, which can improve clustering by increasing between-cluster distances but may also raise within-cluster distances. Consistent with this, WGT performs particularly well on metrics like Dunn and Silhouette, which heavily penalize clustering with insufficient separation. In contrast, CH and *tr*(*W* ^−1^*B*) metrics show more mixed results, with no single transformation clearly outperforming the others.

**FIG 13.**
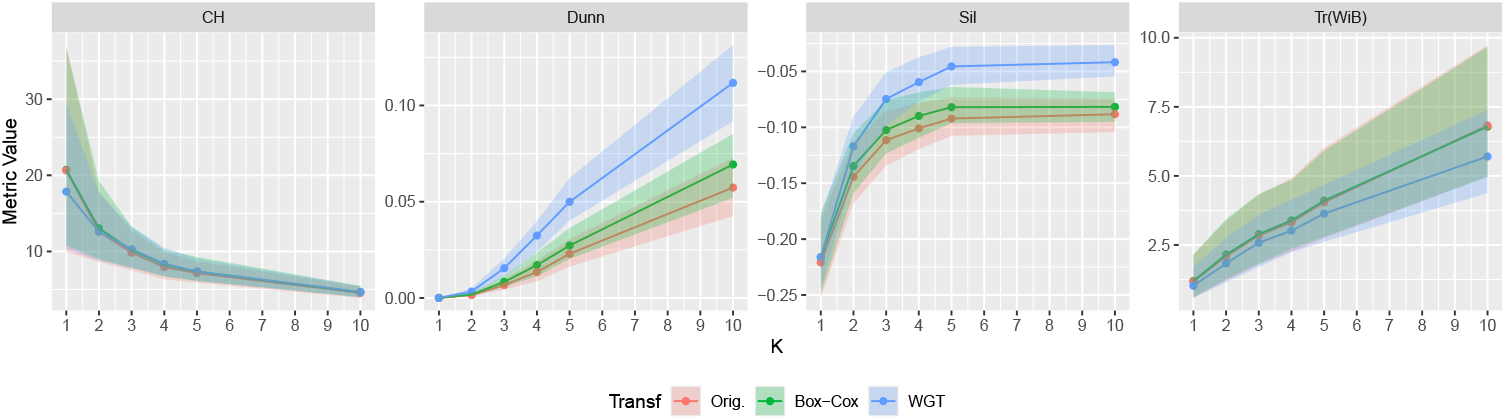
Clustering metrics computed over 100 Monte Carlo subsamples show that WGT generally improves cluster separation by increasing between-cluster distances, with strongest gains seen in Dunn and Silhouette scores.

To quantitatively assess how WGT reduces the variability of analyses such as PCA, we examine the stability of PCA weights under Monte Carlo subsampling. Specifically, we retain all image features and repeatedly subsample 10% of the observations across 100 Monte Carlo replicates. For each subsample, we perform PCA and record the weights assigned to each image feature. We then compute the standard deviation of these weights across subsamples for each principal component. Figure 14 shows box plots of these standard deviations, with one subplot per principal component. The results indicate that while Box-Cox reduces PCA weight variability compared to no transformation, WGT consistently leads to even more stable PCA weights, demonstrating greater robustness across subsamples.

**FIG 14.**
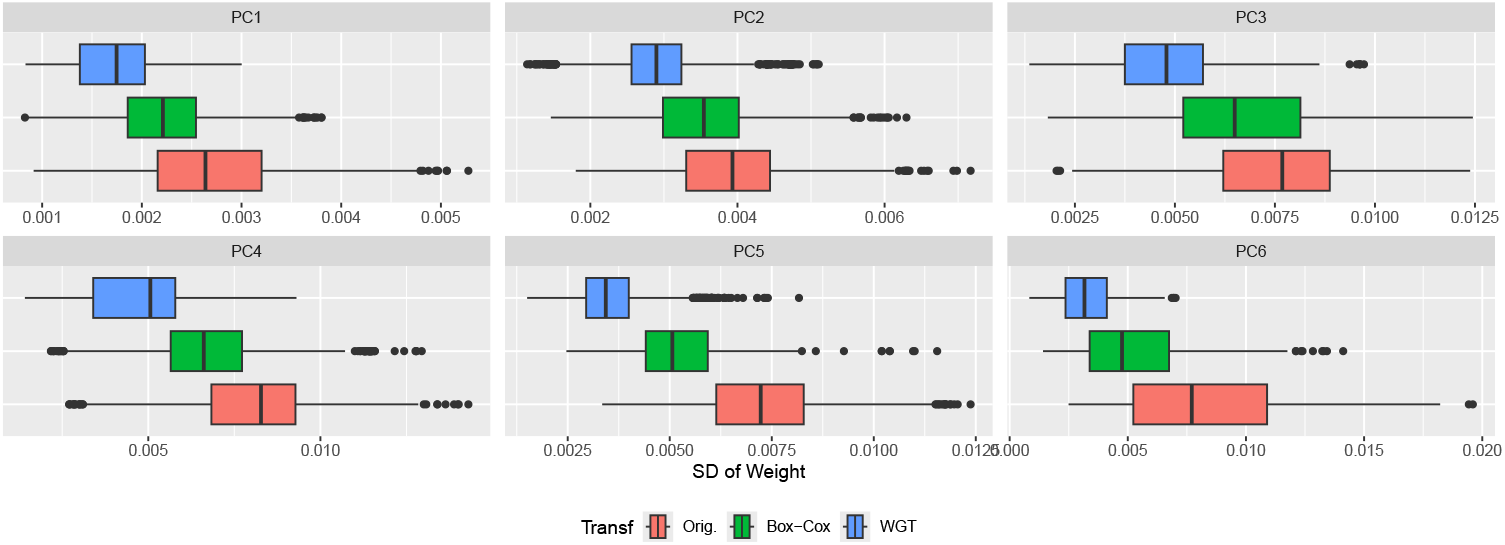
Box plots of PCA weight variability across 100 Monte Carlo subsamples show that WGT consistently produces more stable feature weights than either no transformation or Box-Cox. This suggests that WGT enhances the robustness of PCA to data perturbations.

### 4.4. Identifying Unwanted Spatial Signal

The robustifying effects of WGTs also aid in detecting unwanted spatial artifacts linked to the physical layout of the 384-well plates. Figures 15 and 16 display heatmaps of all sixteen replicate plates across the three transformations. Each subplot reflects the actual spatial arrangement of wells. The subplots themselves are arranged corresponding to replicates (left to right) and to sites (top to bottom).

**FIG 15.**
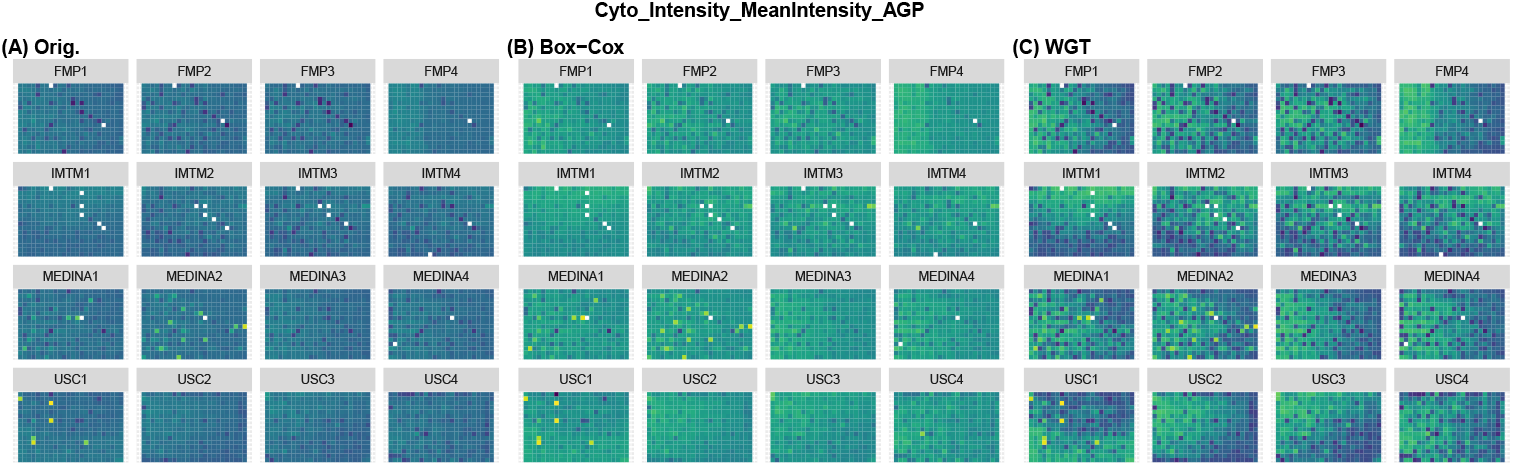
Heatmaps of physical layout of wells across all sixteen replicate plates and the three transformations. Color indicates value of the feature Cyto_Intensity_MeanIntensity_AGP with blue being lower and green being higher. The names correspond as follows: FMP = Leibniz-Forschungsinstitut für Molekulare Pharmakologie; IMTM = Institute of Molecular and Translational Medicine; MEDINA = Fundación MEDINA; USC = Universidad de Santiago de Compostela.

**FIG 16.**
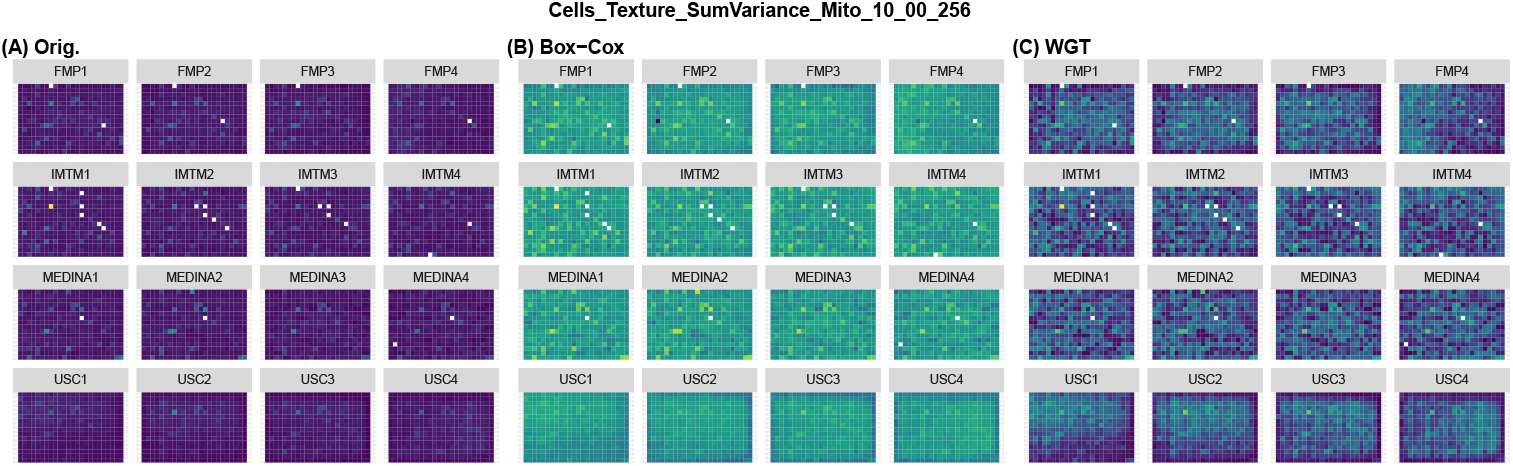
Heatmaps of physical layout of wells across all sixteen replicate plates and the three transformations. Color indicates value of the feature Nuc_Texture_SumVariance_DNA_5_01_256 with blue being lower and green being higher. The names correspond as follows: FMP = Leibniz-Forschungsinstitut für Molekulare Pharmakologie; IMTM = Institute of Molecular and Translational Medicine; MEDINA = Fundación MEDINA; USC = Universidad de Santiago de Compostela.

**FIG 17.**
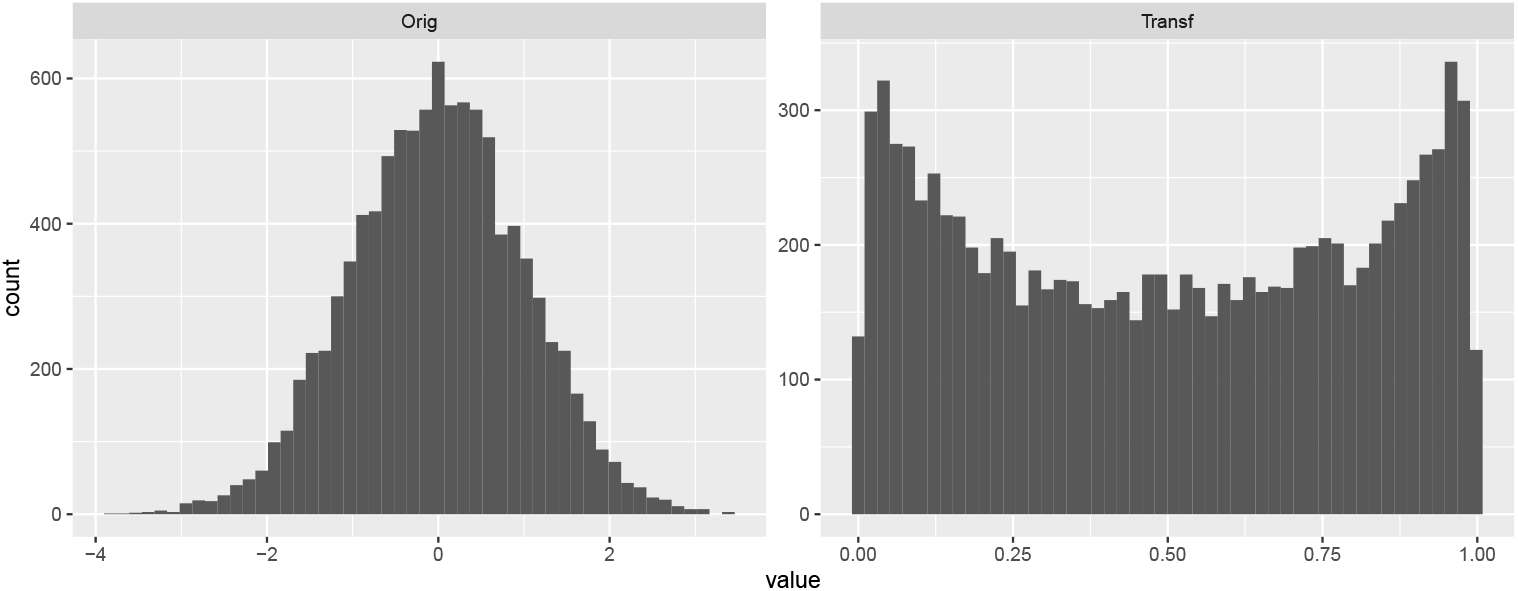
Example of a tail-attenutating transformation. Left panel displays a random sample of N (0, 1) before transformation. Right panel displays the same data after a logistic transformation. After the transformation we now see the data is bimodal since the tails of the original distribution have been accumulated in two piles near the edge of the support by the logistic transformation.

**FIG 18.**
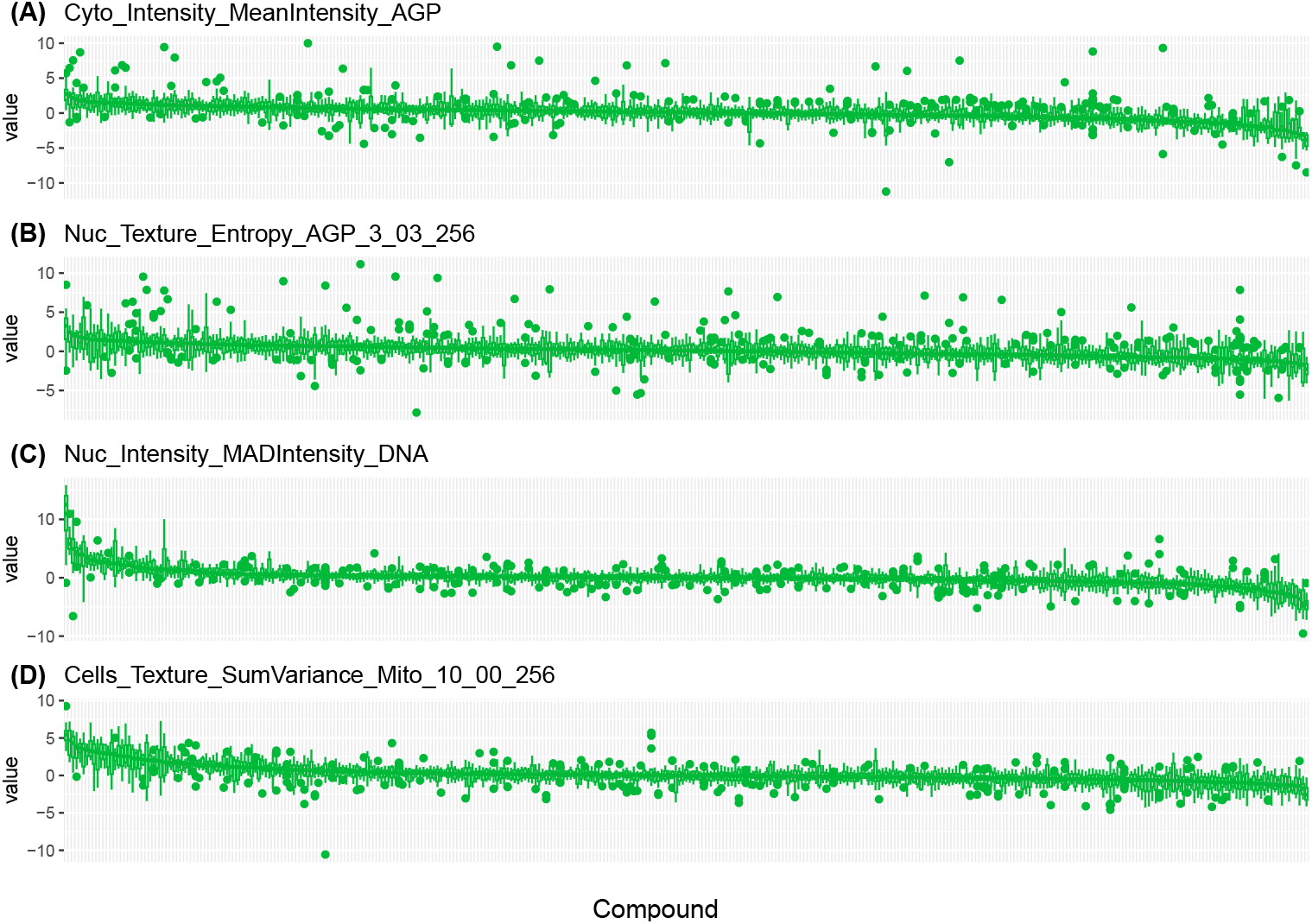
Re-analysis of features in Figure 4 after Box-Cox.

Without transformation (subplots (A)), spatial effects may be hard to discern. This is largely due to the influence of outlying values, which compress the color scale and obscure subtler variation. If our sole interest were in identifying the most extreme wells, this might be acceptable. However, those points are often non-biological, and even when they are not, they can mask other meaningful variation. After applying WGT, striking spatial patterns become visible. In Figure 15, we observe several top-bottom and left-right gradient patterns across wells. In Figure 16, a prominent edge effect appears, with wells along the plate edges showing systematically lower values.

Identifying such spatial effects should be a routine step in high-throughput imaging analysis, as they can significantly impact downstream results. For instance, in this study, all control wells are placed in the two rightmost columns of each plate, which may be problematic in light of the observed edge effects. Spatial visualization can also provide some insight into observations noted in other analyses. For example, the non-biological outliers seen in subplots (A) of Figures 4 and 9 can be examined in light of these plate-wise plots. Firstly, note that the very large values are coming from the first two replicates of MEDINA and first replicate of USC, which may be useful to know when refining experimental procedures. More subtly, in Figure 15 (C), most plates show lower values in the bottom right; however, the USC1 plate (column 1, row 4) deviates from this pattern with unusually high values in that region. This additionally helps account for the large outliers we previously observed. Considering Figure 16, it is worth noting that several earlier outliers lie along the plate edges, which is likely related to the edge effects.

## 5. Conclusion

In this work, we introduced Wasserstein Geodesic Transformations (WGTs), a scalable method for attenuating long tails and outliers in high-throughput cell imaging data. WGTs address a central challenge in such datasets: the presence of highly heterogeneous, skewed, and heavy-tailed image features that distort visualization, calculation of common metrics, and hamper downstream analyses. Unlike traditional approaches, which may be insufficiently adaptive, WGTs provide a general, robust, and tunable transformation approach.

In analysis of real Cell Painting data from Wolff et al. (2025), we found that WGTs substantially reduce outliers, stabilize pairwise correlations, and improve the clarity of data visualizations. They can also help better reveal latent structure when using techniques like PCA or when clustering. In general, we find that WGT increases the robustness of many downstream analyses. These empirical findings support the idea that WGT is valuable for exploring and analyzing complex, high-dimensional imaging datasets. More broadly, WGTs contribute to a growing toolkit for robust, scalable analysis of heterogeneous data. While our focus here was on high-throughput imaging, the formulation and flexibility of WGTs make them readily applicable to other domains with similar challenges where long-tailed, highly variable features are common and problematic.

## Software

An implementation of WGT and tutorial of its use may be found at gjhunt.github.io/wgt.

## Data and Analysis

A docker image and files for conducting the analysis may be found on zenodo at 10.5281/zenodo.14925745.

## 6. Supplementary Material

### *6*.*1. Distancing Tuning*. The CDF corresponding to *g*_*θ*_ is 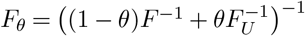. Thus

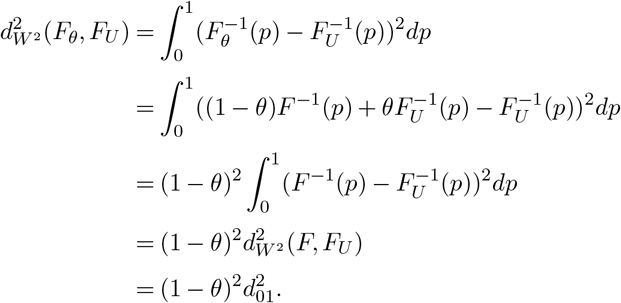

Consequently,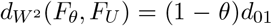. A similar argument shows that 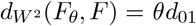.

### 6.2. Density Order Preserving

If our target is (without loss of generality) a *U* (0, 1) distribution, then the Wasserstein geodesic transformation is of the form *g*(*x*) = (1−*θ*)*x* + *θF* (*x*). Any such transformation is density order preserving since *g*^*′*^(*x*) = 1 − *θ* + *θf* (*x*). Thus

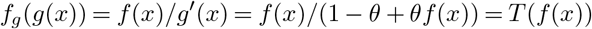

for *T* (*u*) = *u/*(1 − *θ* + *θu*). Here, *T* (*u*) is strictly increasing in *u* for 0 ≤ *θ <* 1 since

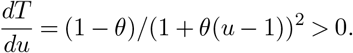

(*dT/du* = 0 when *θ* = 1).

Furthermore, the only Wasserstein geodesic transformations which preserve density order over all *θ* for all possible starting points *F* are those where *F* ^*∗*^ is uniform. To see this, consider the case where *F* is *U* (0, 1). Heuristically, *F* ^*∗*^ has to be uniform since we start as uniform *F* and thus the only way to preserve the uniform density is to transform into another uniform. If we end up at anything non-uniform then we’ve broken DOP at some point. More formally, consider any *y*_1_ and *y*_2_ and let *x*_1_ = *g*^−1^(*y*_1_) and *x*_2_ = *g*^−1^(*y*_2_). Since *F* is uniform then we know that *f* (*x*_1_) = *f* (*x*_2_). Consequently, since *g* is DOP then it must be that *f*_*g*_(*g*(*x*_1_)) = *f*_*g*_(*g*(*x*_2_)) and hence *f*_*g*_(*y*_1_) = *f*_*g*_(*y*_2_). Since this works for any *y*_1_ and *y*_2_ this means that *f*_*g*_ is uniform.

### 6.3. Tail Dominance

Let *F*_*θ*_ be the CDF of *g*_*θ*_(*X*) = (1 − *θ*)*X* + *θF* (*X*) and let 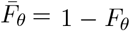 and be the 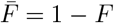 complementary CDFs of *X* and *g*_*θ*_(*X*). We will show two results. First, we will show that for all *θ* ∈[0, 1] the right tail of *X* dominates the right tail of *g*_*θ*_(*X*) such that

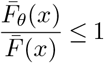

for sufficiently large values of *x*. Secondly, we will show that if (1) *f* is strictly decreasing for large enough *x*, and, (2) that

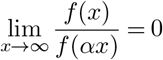

for all *α* ∈ [0, 1] then

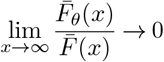

so that the tail of *X* strongly dominates the tail of *g*_*θ*_(*X*). Note that our second conditions is essentially a condition on how fast *f* decreases.

In these cases, 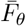 decreases much more quickly than 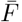. This is true of many common distributions like the Normal, Exponential, LogNormal, or others with more rapidly decreasing tails. Notably, however, it will not be true for distributions with polynomially decreasing tails like the Pareto. While the tail of *X* still dominates the tail of *g*_*θ*_(*X*) for such polynomial-tailed distributions, it does not strongly dominate the tail of *g*_*θ*_(*X*).

#### Proof of dominance

First, note that for sufficiently large *x* we have that *F* (*x*) *< x* and so *g*_*θ*_(*x*) = (1 −*θ*)*x* + *θF* (*x*) *<* (1 −*θ*)*x* + *θx* = *x*. Conversely, for large enough *x* we have that 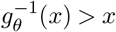. Thus 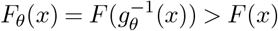 since *F* is increasing. Thus, 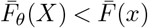 and so *F*_*θ*_(*x*)*/F* (*x*) *<* 1. Thus the tail of *X* dominates the tail of *g*_*θ*_(*X*).

#### Proof of strong dominance

By L’Hôpital’s rule we have that

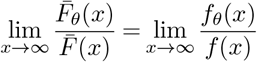

and since *g*_*θ*_(*x*) *→* ∞ as *x →* ∞ then

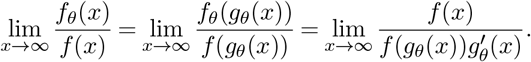

since 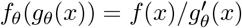. Now since 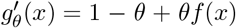 then 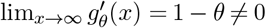 and so

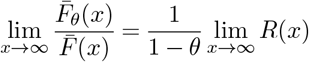

where

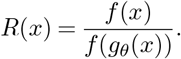

Thus, our goal is to show that *R*(*x*) *→* 0 as *x →* ∞.

For any *α* ∈ [0, 1] if *α >* 1 − *θ* then since *F* (*x*) *<* 1 we have

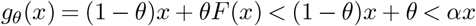

for sufficiently large *x* (in particular, *x > θ/*(*α* + *θ* − 1)). Consequently for sufficiently large *x* we have that

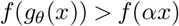

assuming that *f* is strictly decreasing for large enough *x*. This means

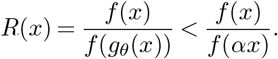

Thus, if the limit of the right-hand quantity is zero as *x* → ∞ for any *α* ∈ [0, 1] then *R*(*x*) → 0 by the squeeze theorem. This completes the result.

#### Commentary

For many common distributions with sufficiently decreasing densities we can show that *f* (*x*)*/f* (*αx*) *→* 0. As examples, consider the standard Gaussian:

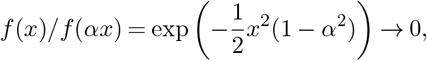

the standard Exponential distribution:

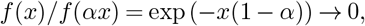

and the standard LogNormal distribution:

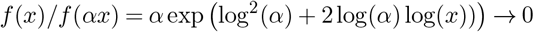

(since log(*α*) *<* 0).

Notably, this will not work for distributions with polynomially decreasing tails. For example, considering the Pareto distribution with fixed minimum of one and shape parameter *β* we find that

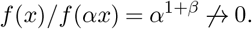

In this particular example, a direct calculation shows that

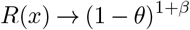

and thus

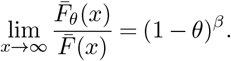

Thus, while the tail of *X* will not strongly dominate the tail of *g*_*θ*_(*X*) we do have that, in the limit, the tail probability of *g*_*θ*_(*X*) will be some small multiple (1 − *θ*)^*β*^ of the tail of probability of *X*. Notably, this multiplier monotonically decreases as *θ* → 1 so that the tail probability of *g*_*θ*_(*X*) is ever smaller multiples of that of *X* as we crank up *θ*.

### 6.4. Linear re-scaling

Note that

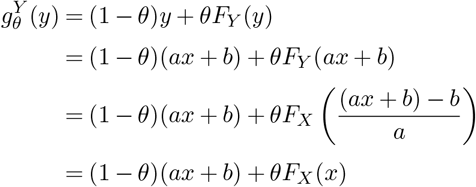

continuing, we find that

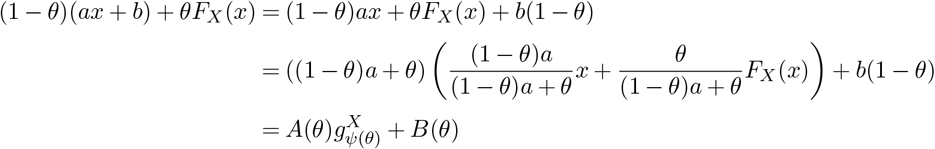

where *ψ*(*θ*) = *θ/*(*a*(1 − *θ*) + *θ*), *A*(*θ*) = *a*(1 − *θ*) + *θ*, and *B*(*θ*) = *b*(1 − *θ*).

#### 6.5. Correlation Tuning

Let *F* = *F*_*X*_(*X*) and for some *θ* let *X*_*θ*_ = (1 − *θ*)*X* + *θF*. Define *J* (*θ*) = Cor (*X*_*θ*_, *F*). Then

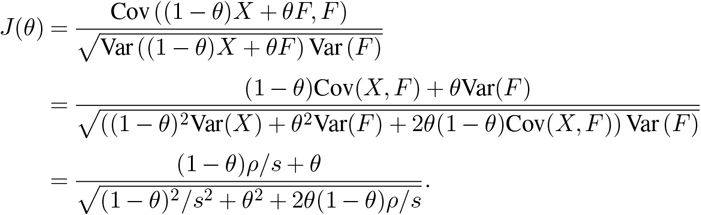

where *ρ* = Cor (*X, F*) and 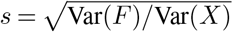. Then one may show that

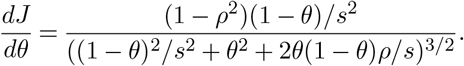

Consequently, *dJ/dθ >* 0 and so *J* is strictly increasing as long as *ρ <* 1 and *θ <* 1.

Now, solving *J* (*θ*) = *ν* is a quadratic equation in *θ*. Consequently, one may apply the quadratic formula. After enough simplification, this formula yields

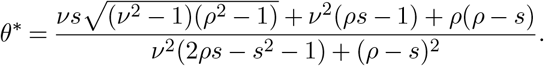

For two random variables *A* and *B*, the squared Wasserstein 2-distance between their distributions can be written as

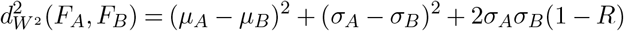

where the *µ*s and *σ*s are the respective means and standard deviations and *R* is the correlation between 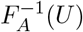 and 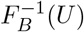 where *U ∼ U* (0, 1). Consequently,

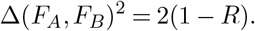

Generally, if *A* ∼*F*_*A*_ and *B* ∼ *F*_*B*_ is is not true that *R* = Cor(*A, B*) since *A* and *B* may have many different correlation values depending on their joint distribution. It is true, however, that *R* is the maximum correlation over all possible joint distribution of *A* and *B*. This maximum correlation is achieved if and only if *A* and *B* are increasing transformations of one another. Consequently, if *A* and *B* are random variables with a joint distribution such that one is an increasing transformation of the other, then it is true that *R* = Cor(*A, B*).

With this in mind, let Cor(*X, F* (*X*)) = *ρ* as previously. Then with *F*_*X*_ as the distribution of *X* we get

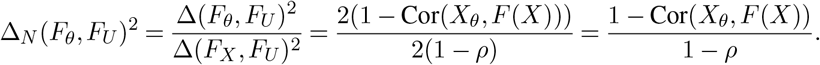

Presuming that Cor(*X*_*θ*_, *F* (*X*)) = *ν* then if we choose *ν* = 1 − (1 − *ρ*)(1 − *η*)^2^ we have

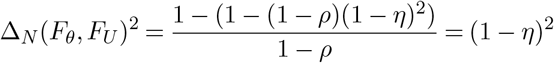

and thus finally we get

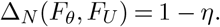

